# More than 90% of nacre matrix proteins are composed of silk-like proteins

**DOI:** 10.1101/2022.09.07.507049

**Authors:** Xiaojun Liu, Zehui Yin, Zhuojun Ma, Jian Liang, Liping Yao, Rongqing Zhang

**Affiliations:** Department of Biotechnology and Biomedicine, Yangtze Delta Region Institute of Tsinghua University, Zhejiang 314006, China; Protein Science laboratory of the Ministry of Education, Tsinghua University, Beijing 100084, China; Taizhou innovation center, Yangtze Delta Region Institute of Tsinghua University,Zhejiang 318000,China; Zhejiang Provincial Key Laboratory of Applied Enzymology, Yangtze Delta Region Institute of Tsinghua University, 705 Yatai Road, Jiaxing 314006, PR China; Chinese academy of fishery sciences, Beijing 100141, China; State Key Laboratory of Plateau Ecology and Agriculture, Qinghai University, Xining 810016, China; Key Laboratory of Freshwater Aquatic Genetic Resources, Shanghai Ocean University, Ministry of Agriculture, Shanghai 201306, China

**Keywords:** nacre, shell matrix proteins, silk-like proteins, proteome

## Abstract

A proteome is a powerful tool to study nacre biomineralization that occurs in an ordered microenvironment composed mainly of proteins and polysaccharides. As more and more proteins are detected, new questions arise about which proteins are responsible for forming this microenvironment. In this study, shell matrix proteins (SMPs) of nacre were analyzed using label-free quantitative proteome. A total of 99.89% of the insoluble nacre SMPs were composed of silk-like proteins, chitin-binding proteins, and cysteine-rich SMPs, which were responsible for organic framework assembly. A total of 99.34% of the soluble nacre SMPs were composed of silk-like proteins and chitin-binding proteins, which were responsible for forming protein gel filling in organic frameworks. The content of silk-like proteins was more than 90% in both insoluble and soluble nacre SMPs. As organic frameworks and protein gel together constructed a microenvironment for calcium carbonate biomineralization, these results provided a novel understanding of nacre formation.

## Introduction

The beautiful luster of nacre in shells and pearls has been loved by people since ancient times, and the increased interest of materials scientists today is because of its excellent material characteristics, such as high fracture resistance^1^. These excellent properties of nacre originate from its microstructure in which nanoscale aragonite platelets are arranged in continuous parallel layers and separated by interlamellar and intercrystalline organic sheets^2-4^. What gives shellfish such a highly ordered self-assembling ability *in vitro* has long been the interest of scientists. The chemical composition of nacre naturally has become the primary concern of scientists in solving this problem. Nacre typically consists of more than 95% calcium carbonate and up to 5% organic matrix, which is composed of shell matrix proteins (SMPs), polysaccharides, lipids, and so on. Of all the organic matrices, SMPs are the most studied, in part because of the ease of the study of proteins and advances in protein research technology. For example, many polysaccharide components are derived from protein modification except chitin^5^, making the separation of polysaccharide components a complex and difficult task.

From the 1950s to the 2000s, many proteins, such as Nacrein, MSI60, and MSI31 in *Pinctada fucata*^6,7^, were successfully identified using the protein and gene technologies, such as gel separation, high-performance liquid chromatography, and molecular biology techniques^8^. Then, the shell proteomes, which were named “shellome” by Marin^9^, gave us the “biomineralization toolkit” to understand the biomineralization role of SMP as a whole^10^. Nacre proteomic analysis was conducted in *Nautilus macromphalus*^11^, *Haliotis asinina*^12^, *Unio pictorum*^13^, *Pinctada margaritifera* ^14^, and the edible mussel *Mytilus*^15^. Then, this powerful “biomineralization toolkit” first provided the evidence that Prisms and nacre were assembled from very different protein repertoires in *P. margaritifera* and *Pinctada maxima*, and these layers did not derive from each other^10^. After that, more nacre proteomes were reported, such as in *H. asinina*^16^, *Hyriopsis cumingii*^17^, *P. fucata* ^18,19^, *Elliptio complanata*^20^, *Villosa lienosa*^20^, and *Haliotis laevigata*^21^.

Most of these shell proteomes were qualitatively investigated, and the abundance of SMPs was not considered^9^. Mann et al. first identified major proteins in nacre and prismatic organic shell matrix based on the abundance using proteomic and next-generation sequencing was true quantitative proteomics^21^. This study was the most comprehensive for nacre SMPs in *Haliotis* species, and quantitative proteomics proved to be a powerful analytical tool in the study of SMPs. MSI60-related proteins mainly existed in nacre-insoluble SMPs, but the percentages of MSI60-related proteins (4.38%–8.64%) were not very consistent with previous studies^21^. X-ray diffraction studies showed that nacre-insoluble organic frameworks were mainly composed of silk-fibroin-like proteins^22,23^. The amino acid composition analysis in previous studies showed that Glycine, Alanine, Serine, and Aspartic acid were richer than other amino acids in the nacreous layer^24,25^. This might mean that some proteins and protein families were also richer than many other proteins in nacre SMPs. The high content of glycine, alanine, and serine was the main characteristic of silk-like proteins^7^. Synthetic nacre exhibiting good ultimate strength and fracture toughness was successfully synthesized with a matrix composed of layered chitin and filled silk fibroin predesigned according to the nacre formation model^26,27^. This also proved the importance of silk-like proteins in nacre biomineralization. Therefore, a new question arose about whether silk-like proteins were the main components of nacre. Nacre SMPs of *H. cumingii* were studied using label-free quantitative proteome to solve this issue.

## Results

### 1. Protein composition of nacre SMPs

A total of 426 and 345 peptides were detected in nacre EDTA-soluble SMPs (ESMs) and nacre EDTA-insoluble SMPs (EISMs), respectively, belonging to 133 and 95 proteins (figure S1-4). Most identified peptides matched the expressed sequence tags (ESTs) sequences of *H. cumingii*. Ten known SMPs were found in the nacre proteome, including hic74^28^, silkmapin^29^, silkmaxin^30^, hic9^31^, HcKuPI^32^, hic7^33^, hic14^34^, hichin^35^, theromacin^36^, and an unpublished protein hc-gastric intrinsic factor-like protein 2 (GIFLP2) (Table S1-2). And we cloned the completed cDNA sequences of these unknown SMPs based on ESTs (Table S1-2). The completed cDNA sequences of 55 unknown SMPs were successfully cloned from mantle tissue (figure S5-9), and only 8 peptides (Feb-85, Jan-00, Feb-89, Jan-28, 1036-1, Jan-67, Jan-33, and 386-1) could not be cloned. In this process, some different peptides from different sequence regions of a protein were successfully classified. In the end, a total of 73 proteins were found in the nacre proteome (Table 1-2), and 66 and 48 proteins were found in ESMs and EISMs, respectively (Figure 1). Among them, 41 proteins existed in both ESM and EISM. The characterization of the full sequence of 89% SMPs allowed us to obtain an overall map of nacre SMPs.

**Table 1.**
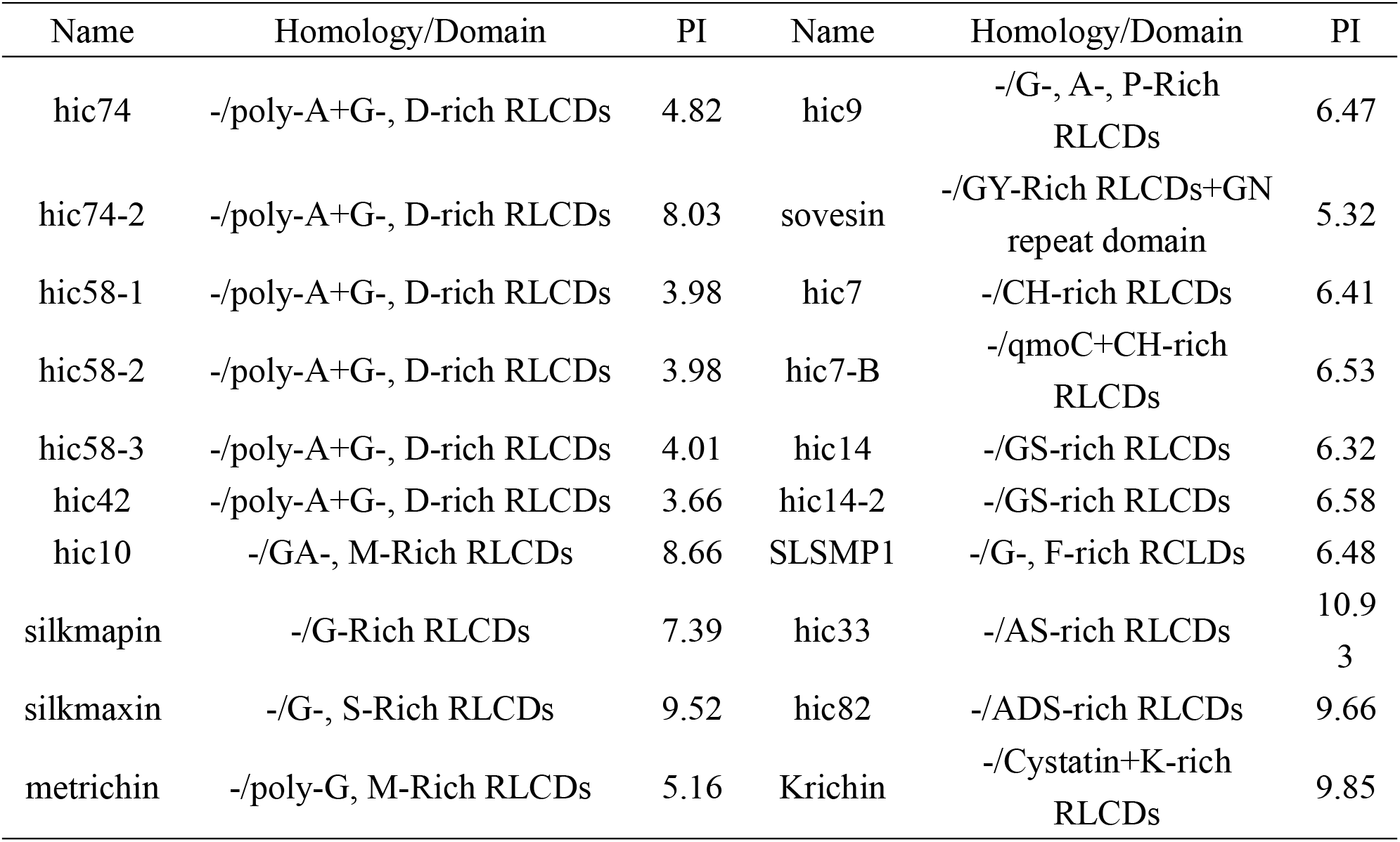
RLCDs/LCDs containing matrix proteins with completed amino acid sequence

**Table 2.** Other matrix proteins with completed amino acid sequence

| Name | Homology/Domain | PI | Name | Homology/Domain | PI |
| --- | --- | --- | --- | --- | --- |
| hic124 | -/2ChtBD2+LamG | 5.16 | upsalin2 | Upsalin *Upi*- | 6.52 |
| hichin | -/ChtBD2 | 4.59 | upsalin3 | Upsalin *Upi*- | 8.49 |
| hic54 | -/2ChtBD2+KAR9 | 6.45 | upsalin4 | Upsalin *Upi*- | 8.1 |
| hc-pif2 | Pif *Pfu*/vWA+2chitin-binding domains | 4.46 | cysrichin-A | -/- | 8.02 |
| hic92 | COL6A<br>*Mco*/LamG+ChtBD2+vWA | 6.63 | cysrichin-B | -/- | 8.52 |
| chitinase-4 | Chitinase *Mga*<br>*Hcu*/ChiA+2ChtBD2 | 6.92 | cysrichin-C | -/- | 7.79 |
| prismafin | Kunitz/Bovine pancreatic trypsin inhibitor domain protein *Tsu*/2KU | 7.91 | cysrichin-D | -/WAP | 7.59 |
| hic118 | Hexosaminidase *Mga*/ Med15+<br>2CHB_HEX+GH20 | 6.23 | cysrichin-E | -/- | 7.51 |
| hic68 | collagen-like protein<br>*Eco*/collagen+PRK07764+PRK149<br>59 | 8.51 | tofesin | -/- | 8 |
| hc-NPAC | nascent polypeptide-associated complex subunit alpha *Cgi*/NAC | 4.67 | hic6 | -/- | 4.01 |
| hc-RBL13 | 60S ribosomal protein L13-like<br>*Cvi*/ Ribosomal_L13e | 11.3 | hc-beta-actin2 | Actin *Dch*/actin | 5.39 |
| hc-GTLL | glycine--tRNA ligase-like<br>*Pca*/PLN02734 | 7.85 | hic5 | -/- | 5.22 |
| hc-EF1A | translation elongation factor<br>1-alpha *Pli*/SR rich | 10.6<br>6 | argrichin | -/- | 8.3 |
| hc-ETIF3 | eukaryotic translation initiation factor 3 subunit G-like<br>*Cgi*/eIF3G+RRM | 8.43 | sonefin | -/- | 8.51 |
| zonesin | Mucin-2 like *Nve*- | 3.7 | teosin2 | Teosin *Hcu*- | 7.66 |
| hc-RBPL | riboflavin-binding protein-like<br>*Lan*/Folate_rec | 4.86 | fiverin | -/- | 4.78 |
| hc-MCSGL | mesenchyme-specific cell surface glycoprotein-like *Mye*- | 4.91 | hic8 | -/- | 9.38 |
| hc-ZFP385<br>B | zinc finger protein 385B *Lan* /<br>ZnF_U1+ zf-met+zf-C2H2_jaz | 8.93 | hc-vdg3 | vdg3 *Med*- | 6.51 |
| hc-CDIC | chitin deacetylase *Gae*/PRK10246<br>+TolA+CE4_CDA_like_2 | 9.97 | hic15 | -/2WAP | 6.97 |
| hc-GIFLP2 | gastric intrinsic factor-like *Pca*<br>/DUF4430 | 8.82 | hic16 | -/- | 9.18 |
| hic51 | -/antiphage_ZorA_3 | 6.48 |  |  |  |

**Figure 1.**
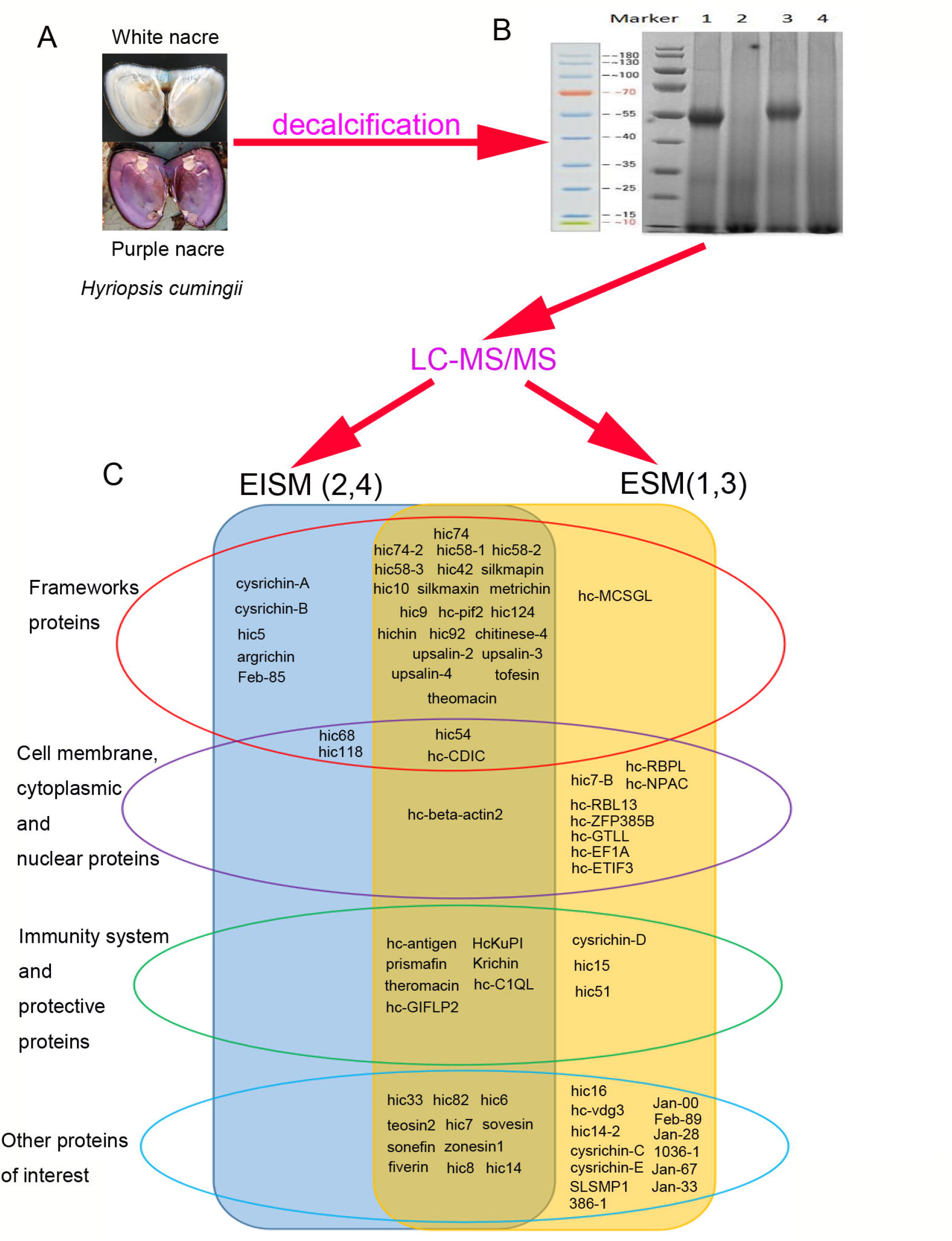
Nacre label-free proteome analysis and the proteins detected in the proteome. (A) Inner surface of shells shows the white and purple nacre. (B) SDS-PAGE analysis of the EISMs and ESMs extracted by nacre powder decalcification. 1, EISMs from white nacre; 2, ESMs from white nacre; 3, EISMs from purple nacre; 4, ESMs from purple nacre. (C) Proteins detected in the proteome. Proteins in the blue rounded rectangle existed in EISMs. Proteins in the orange rounded rectangle existed in ESMs. Proteins in the red border oval belonged to framework proteins. Proteins in the red border oval belonged to the cell membrane, cytoplasmic, and nuclear proteins. Proteins in the green border oval belonged to the immunity system and protective proteins. Proteins in the light blue border oval belonged to other proteins of interest. A total of 73 proteins were found in the nacre proteome and 66 and 48 proteins in ESMs and EISMs, respectively. Among them, 41 proteins existed in both ESM and EISM. The full sequence of 89% SMPs was successfully characterized except eight proteins (Feb-85, Jan-00, Feb-89, Jan-28, 1036-1, Jan-67, Jan-33, and 386-1).

1. Framework proteins included silk-like proteins, chitin-binding proteins, chitin-related enzymes, extracellular matrix (ECM)-related proteins, and cysteine-rich proteins (Table 1-2). The most common framework proteins were silk-like proteins. In this study, six spider silk-like proteins (hic74, hic74-2, hic58-1, hic58-2, hic58-3, and hic42) and five flagelliform silk-like proteins (silkmapin, silkmaxin, hic10, hic9, and metrichin) were found (Table 1). Spider silk-like proteins (hic74, hic74-2, hic58-1, hic58-2, hic58-3, and hic42) were similar to MSI60^7^ and flagelliform silk-like proteins, silkmapin, and silkmaxin were similar to the members of shematrin family^37^; however, all similarity existed in the primary structure of proteins with no homology. The hic58 family and hic42 were the second and third kinds of spider silk-like proteins in *H. cumingii*. The hic58 family contained three members, and hic74-2 was a new member of the hic74 family. Therefore, the alternative splicing seemed to exist in the hic74 family and hic58 family according to different protein lengths (Figure S10-11). Silk-like proteins had typical repetitive low complexity domains (RLCDs) (Table 1). Comprising one or more amino acid–rich RLCDs was the main characteristic of many important SMPs^38^. However, the role of these RLCDs in biomineralization remains unknown. RLCDs also could be found in some framework proteins such as hic14, hic14-2, and SLSMP1 (Table 1).
Chitin-binding domains were found in six proteins in the nacre proteome, hichin, hc-pif2, hic124, hic92, hic54, and chitinase-4 (table 2). hc-pif2 had eight amino acids mutation with pif previously reported in *H. cumingii*^39^, and a “DIYKRKDRDYQS” amino acids sequence was missing in hc-pif2 (Figure S12). Chitinase-4 comprised a chitin-binding domain probably because it played a role in the chitin framework modification. Another chitin-related enzyme was hc-CDIC (chitin deacetylase isoform C) containing the CE4_CDA_like_2 domain (putative catalytic domain of chitin deacetylase). The ECM-related protein hic118 comprised domains that interacted with the saccharide moieties: two CHB_HEX domains (putative carbohydrate-binding domain), a GH20 domain, and a Med15 domain. GH20 was a glycosyl hydrolase family 20 (GH20) domain of the beta-*N*-1,4-acetylhexosaminidase and could hydrolyze the beta-1,4-glycosidic linkages in oligomers derived from chitin in *Serratia marcescens*^40^. The chitin-binding proteins also comprised multidomains except for hichin (Table 2). These domains were mainly typical ECM-related domains, such as the laminin_G domain in hic124 and hic92 and von Willebrand type A domain in hc-pif2 and hic92. Other ECM-related proteins include mucin-like protein zonesin, collagen-like protein hic68, and mesenchyme-specific cell surface glycoprotein-like protein hc-MCSGL. Another kind of important framework proteins, cysteine-rich proteins, had many members in this study, including upsalin2, upsalin3, upsalin4, tofesin, cysrichin-A, cysrichin-B, hic6, teosin2, hic7, prismafin, cysrichin-C, cysrichin-D, cysrichin-E, hic15, and theromacin. However, only upsalin2, upsalin3, upsalin4, tofesin, cysrichin-A, cysrichin-B, and theromacin were framework proteins, as their peptides were detected in the EISM proteome. Other framework proteins included hic5, argrichin, and Feb-85 (Table 2).
2. Cell membrane, cytoplasmic, and nuclear proteins included PRK07764 domain (hic68), PRK14959 domain (hic68), Med15 domain (hic118), and ZnF_U1 domain (hc-ZFP385B) containing nuclear proteins, qmoC domain (hic7-B), TolA domain (hc-CDIC), and Folate_rec domain (hc-RBPL) containing cell membrane proteins and cytoplasmic proteins such as hc-NPAC (nascent polypeptide-associated complex subunit alpha, NAC domain containing), hc-RBL13 (60S ribosomal protein L13-like, ribosomal_L13e domain containing), hc-GTLL (glycine-tRNA ligase-like, PLN02734 domain containing), hc-EF1A (translation elongation factor 1-alpha), hc-ETIF3 (eukaryotic translation initiation factor 3 subunit G-like, eIF3G, and RNA recognition motif (RRM) domains containing), hc-beta-actin2 (actin domain), and KAR9 domain-containing protein hic54 (Figure 1). QmoC (quinone-interacting membrane-bound oxidoreductase complex subunit), Folate_rec (folate receptor family domain), TolA (membrane protein involved in colicin uptake), Med15 (transcription activators domain), NAC, PLN02734 (glycyl-tRNA synthetase), eIF3G (eukaryotic translation initiation factor 3 subunit G), RRM, KAR9, PRK07764, and PRK14959 (DNA polymerase III subunits gamma and tau domains) domains were first found in nacre SMPs. KAR9 domain existed in cytoskeletal proteins required for karyogamy, correct positioning of the mitotic spindle, and orientation of cytoplasmic microtubules in the yeast *Saccharomyces cerevisiae*^41^.
3. Immunity system and protective proteins included Kunitz (KU) domain (hc-antigen, HcKuPI, and prismafin), cystatin domain (Krichin), WAP domain-containing proteolytic protective proteins (cysrichin-D and hic15), Macin domain (theromacin), antiphage_ZorA_3 domain (hic51), and C1q domain (hc-C1QL) containing immune system proteins, and hc-GIFLP2 (gastric intrinsic factor-like protein) (Figure 1). Cystatin (cysteine protease inhibitors) and antiphage_ZorA_3 domain (antiphage defense ZorAB system ZorA) domains were first found in nacre SMPs. Proteins of antiphage_ZorA_3 domain were putative H^+^ channel proteins and were reported to be involved in the antiphage defense^42^.
4. Other proteins of interest were listed in Figure 1. Besides G-or A-rich RLCDs mentioned above, hic7 and hic7-B containing CH-rich RLCDs. Except for RLCDs, the repeats of hic33 and hic82 sequences were extremely high, accounting for 97.9% and 100% of mature protein sequences, respectively. hic33 consisted of 22 repetitive “(D/E/N)(A/L/V/I)PASFVSLSKRS(A/T)G,” and hic83 consisted of 53 repetitive “(D/N)APASFISLSKRS(A/T)(E/D/G/A)” (figure S13). Enzymes were not listed separately because fewer enzymes were found in this study than in other studies, and the import enzymes in other shell proteomes, such as carbonic anhydrase and tyrosinase, were not found in this study^8,9^. Only chitinase-4, hc-CDIC, hexosaminidase (hic118), and hc-GTLL were found in this study.

Some proteins contained multiple domains whose biological functions differed widely, resulting in their classification as different proteins. hic68, hic118, hc-CDIC, and hic54 contained the domains belonging to ECM (collagen domain in hic68; CE4_CDA_like_2 domain in hc-CDIC; ChtBD2 in hic54; CHB_HEX; and GH20 in hic118) and the domains belonging to cytoplasmic and nuclear proteins (PRK07764 and PRK14959 in hic68; Med15 in hic118; KAR9 in hic54; and exonuclease subunit PRK10246 domain and TolA domain in hc-CDIC) (Figure 1; Table 2); therefore, they might also belong to cytoplasmic and nuclear proteins. The differences of some protein family in cDNA and amino acid sequences, such as hic7 family, hic14 family, teosin family and upsaline family, were show in figure S14.

### 2. Amino acid composition of SMPs with full-length sequence

The amino acid composition of 65 nacre SMPs is shown in figure 2. Glycine, alanine, cysteine, proline, and serine are abundant in many proteins. Further, 28 proteins had glycine content greater than 10% and 11 proteins had glycine content greater than 20%. Fourteen proteins contained more than 10% alanine, and six of them contained more than 20% alanine. Sixteen proteins contained more than 10% cysteine. The proline content of 15 proteins was more than 10%. Twenty proteins contained more than 10% serine. Six proteins had arginine content greater than 10%. Five proteins had lysine, leucine, and aspartic acid content of more than 10%. Four proteins contained more than 10% tyrosine.

**Figure 2.**
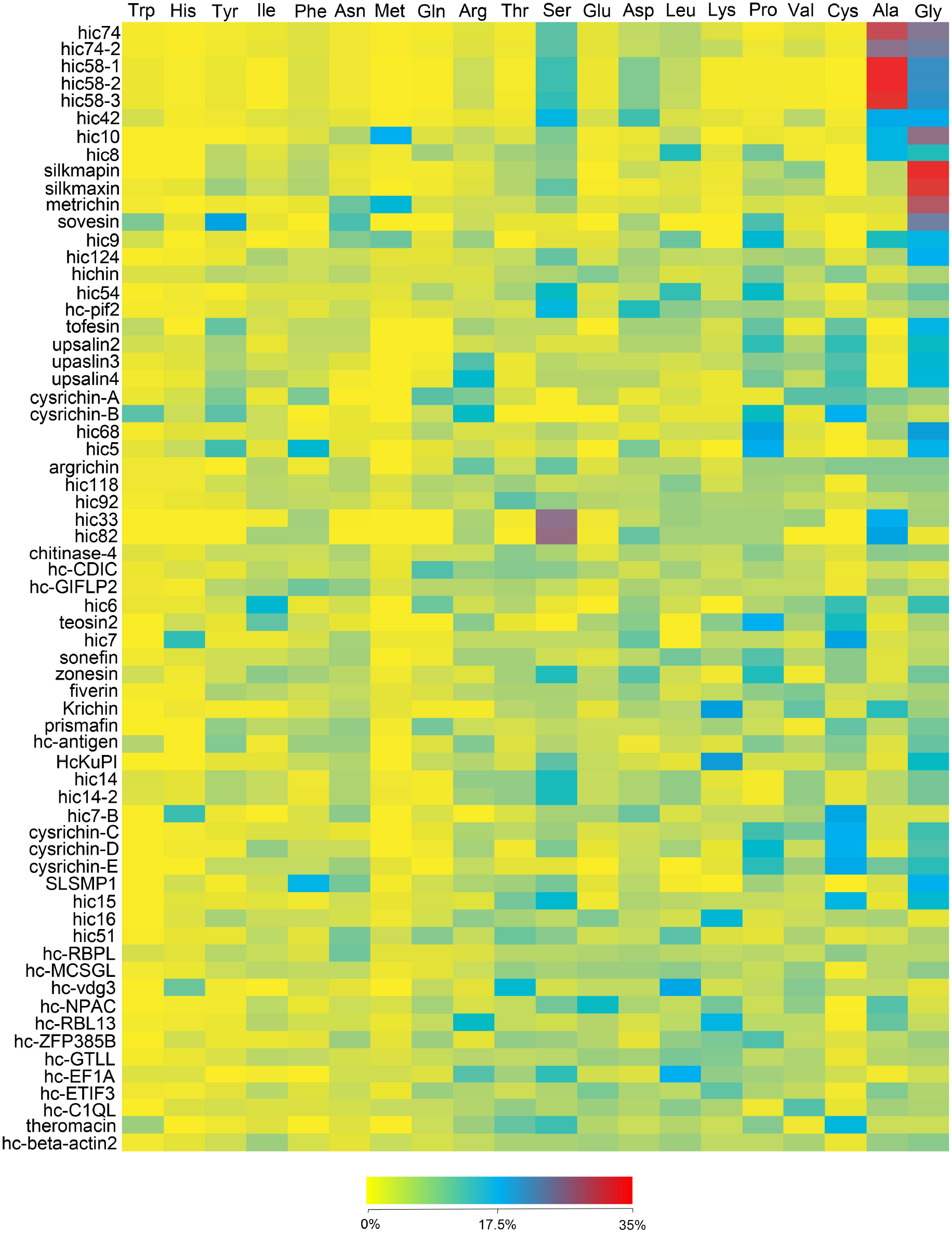
Amino acid composition of 65 SMPs with full-length sequence. All 20 amino acids could be found with percentages of more than 10% in one or some proteins. Nine proteins contained no amino acids more than 10% (hichin, hic118, chitinase-4, fiverin, hc-GIFLP2, hc-RBPL, hc-MCSGL, hc-GTLL, and hc-beta-actin2). Glycine, alanine, cysteine, proline, and serine were abundant in many proteins. Also, only the contents of glycine, alanine, and cysteine were found to be more than 20% in one or proteins.

The number of proteins containing more than 10% of both valine and threonine was 3 (table 3). The number of proteins containing more than 10% glutamine, methionine, phenylalanine, isoleucine, and histidine was 2. The number of proteins with glutamic acid and tryptophan content of more than 10% was 1. Although the content of all 20 amino acids was more than 10% in one or some proteins, 9 proteins contained no more than 10% amino acids (hichin, hic118, chitinase-4, fiverin, hc-GIFLP2, hc-RBPL, hc-MCSGL, hc-GTLL, and hc-beta-actin2). Other proteins contained more than 10% one to five amino acids (Table 3). Twenty-seven proteins contained three amino acids which were more than 10% in their amino acid composition. The total percentage of these amino acids, which was more than 10%, was more than 40% in 23 proteins (Table 3).

**Table 3.** The number, kinds and percent of amino acids more than 10% in protein with full length sequence

| Protein name | Number | Amino acid | Total percent | Protein name | Number | Amino acid | Total percent |
| --- | --- | --- | --- | --- | --- | --- | --- |
| hic74 | 3 | G,A,S | 67 | hc-CDIC | 1 | Q | 11 |
| hic74-2 | 3 | G,A,S | 61.4 | hic6 | 3 | G,C,I | 36.5 |
| hic58-1 | 3 | G,A,S | 68.4 | teosin2 | 3 | C,P,I | 39.6 |
| hic58-2 | 3 | G,A,S | 68.5 | hic7 | 3 | C,D,H | 41.7 |
| hic58-3 | 3 | G,A,S | 67.6 | sonefin | 1 | P | 10.8 |
| hic42 | 4 | G,A,D,S | 63.4 | zonesin | 3 | P,D,S | 35.2 |
| hic10 | 3 | G,A,M | 57.9 | Krichin | 3 | A,V,K | 42.8 |
| hic8 | 3 | G,A,L | 39.3 | prismafin | 1 | C | 10.3 |
| silkmapin | 1 | G | 34.4 | hc-antigen | 1 | G | 10.1 |
| silkmaxin | 2 | G,S | 43.4 | HcKuPI | 3 | G,K,S | 44.2 |
| metrichin | 2 | G,M | 43.3 | hic14 | 1 | S | 12.3 |
| sovesin | 4 | G,P,N,Y | 66.6 | hic14-2 | 1 | S | 12.3 |
| hic9 | 3 | G,A,P | 40.8 | Hic7-B | 2 | C,H | 31.2 |
| hic124 | 2 | G,S | 27.3 | cysrichin-C | 3 | G,C,P | 39.1 |
| hic54 | 3 | P,L,S | 37.2 | cysrichin-D | 3 | G,C,P | 41.8 |
| hc-pif2 | 2 | D,S | 26.5 | cysrichin-E | 3 | G,C,P | 42.8 |
| tofesin | 4 | G,C,P,Y | 45.9 | SLSMP1 | 2 | G,F | 31.8 |
| upsalin2 | 3 | G,C,P | 36.3 | hic15 | 3 | G,C,S | 41.6 |
| upsalin3 | 3 | G,C,R | 35.8 | hic16 | 1 | K | 13.9 |
| upsalin4 | 3 | G,C,R | 38.7 | hic51 | 1 | L | 10.6 |
| cysrichin-A | 3 | C,V,Q | 32.1 | hc-vdg3 | 2 | L,T | 32.9 |
| cysrichin-B | 5 | C,P,R,Y,W | 63.8 | hc-NPAC | 2 | A,E | 23.5 |
| hic68 | 2 | G,P | 41.4 | hc-RBL13 | 3 | A,K,R | 37.5 |
| hic5 | 4 | G,P,F,Y | 59.1 | hc-ZFP385B | 1 | P | 11 |
| argrichin | 2 | S,R | 20 | hc-EF1A | 3 | L,S,R | 39.3 |
| hic92 | 1 | T | 10.6 | hc-ETIF3 | 1 | K | 10.4 |
| hic32 | 2 | A,S | 44 | hc-C1QL | 1 | V | 10.9 |
| hic82 | 3 | A,D,S | 56.9 | theromacin | 3 | C,S,T | 35.7 |

### 3. Quantitative analysis of different protein groups in nacre SMPs

Previous studies revealed that the logarithm of the concentration and the logarithm of the detected peptide number of a protein in proteome had a linear correlation^43-45^. We calculated the percentage of the SMPs in this nacre proteome based on their relative content calculated by the method developed by States et al^45^. The content of most dominant proteins hic74 and hic58-3 was 66.1% and 12.96%, respectively, in nacre EISMs, and 59.21% and 27.53% in nacre ESMs (Table 4). Considering that both proteins are silk-like proteins, the content of all silk-like proteins was calculated. The content of total silk-like proteins was 91.7% and 97.11% in nacre EISMs and ESMs, respectively (Figure 3). Flagelliform silk-like proteins (silkmapin, silkmaxin, hic10, hic9, and metrichin) mainly existed in nacre EISMs (10.78%), and their content was less in ESMs (0.33%) (Figure 3). The content of spider silk-like proteins (hic58 family, hic74 family, and hic42) was 80.92% and 96.78% in nacre EISMs and ESMs, respectively (Figure 3). In conclusion, silk-like proteins, especially spider silk-like proteins, were dominant in nacre SMPs. ECM-related domains containing SMPs were the second dominant kind of SMPs in total nacre SMPs. Their content in ESMs (2.24%) was more than that in EISMs (1.59%). Chitin-binding proteins were dominant in ECM-related domains containing SMPs. Their content were 2.23% and 1.52% in ESMs and EISMs respectively (Figure 3). Cysteine-rich SMPs were also abundant. However, they were mainly found in EISMs (6.66%) and only 0.49% in ESMs (Figure 3). Among them, cysrichin-C, cysrichin-D, cysrichin-E, and hic15 were all only found in ESMs, implying that they were not organic framework proteins. Theromacin and prismafin were both found in ESMs and EISMs. They might play multiple roles in nacre biomineralization: as immunity system and protective proteins and as organic framework proteins. The content of other cysteine-rich proteins (upsalin2, upsalin3, upsalin4, tofesin, cysrichin-A, cysrichin-B, hic6, teosin2, and hic7) was higher in ESIMs. Upsalin family (upsalin2, upsalin3, and upsalin4) (5.37%) and tofesin (1.1%) had higher contents in ESIMs than other cysteine-rich proteins (0.19%). Upsalin was first found in the freshwater mussel *U. pictorum*^46^ and was also characterized as a nacreous layer protein in *H. cumingii*^47^.

**Table 4.** Percent of SMPs in nacre EISMs, ESMs and total SMPs

| Protein name | Percent in EISMs (%) | Percent in ESMs (%) | Protein name | Percent in EISMs (%) | Percent in ESMs (%) |
| --- | --- | --- | --- | --- | --- |
| hic74 | 66.10043 | 59.21335 | sonefin | 0.00097 | 0.00034 |
| hic74-2 | 0.01977 | 0.06989 | zonesin1 | 0.00097 | 0.00034 |
| hic58-1 | 0.88218 | 0.79467 | fiverin | 0.00097 | 0.00034 |
| hic58-2 | 0.08015 | 5.74669 | Krichin | 0.00651 | 0.02780 |
| hic58-3 | 12.95965 | 27.52683 | hic8 | 0.00651 | 0.00034 |
| hic42 | 0.88218 | 3.43292 | hc-antigen | 0.00097 | 0.00034 |
| hic10 | 4.64264 | 0.06989 | HcKuPI | 0.00097 | 0.00686 |
| silkmapin | 3.57569 | 0.01509 | hic14 | 0.00097 | 0.00034 |
| silkmaxin | 0.40110 | 0.00226 | hic14-2 | 0 | 0.00034 |
| metrichin | 0.53533 | 0.24112 | hic7-B | 0 | 0.00034 |
| hic9 | 1.62578 | 0.00034 | cysrichin-C | 0 | 0.00226 |
| hic124 | 1.09849 | 1.81909 | cysrichin-D | 0 | 0.00034 |
| hichin | 0.00651 | 0.10077 | cysrichin-E | 0 | 0.00034 |
| hic54 | 0.08015 | 0.00686 | SLSMP1 | 0 | 0.06989 |
| hc-pif2 | 0.29048 | 0.30602 | hic15 | 0 | 0.00686 |
| hic92 | 0.00097 | 0.00034 | hic16 | 0 | 0.00226 |
| chitinase-4 | 0.04349 | 0.00034 | hic51 | 0 | 0.00034 |
| upsalin2 | 0.88218 | 0.02780 | hc-RBPL | 0 | 0.00034 |
| upsalin3 | 0.40110 | 0.02780 | hc-MCSGL | 0 | 0.00226 |
| upsalin4 | 4.08708 | 0.38106 | hc-vdg3 | 0 | 0.00034 |
| tofesin | 1.09849 | 0.01509 | hc-NPAC | 0 | 0.00686 |
| cysrichin-A | 0.01977 | 0 | hc-RBL13 | 0 | 0.00034 |
| cysrichin-B | 0.00097 | 0 | hc-ZFP385B | 0 | 0.00034 |
| hic68 | 0.01977 | 0 | hc-GTLL | 0 | 0.00034 |
| hic5 | 0.00651 | 0 | hc-EF1A | 0 | 0.00034 |
| argrichin | 0.00097 | 0 | hc-ETIF3 | 0 | 0.00034 |
| hic118 | 0.00651 | 0 | Jan-00 | 0 | 0.00034 |
| Feb-85 | 0.00097 | 0 | Feb-89 | 0 | 0.00034 |
| hic33 | 0.00651 | 0.00226 | Jan-28 | 0 | 0.00034 |
| hic82 | 0.00651 | 0.00034 | 1036-1 | 0 | 0.00686 |
| hc-CDIC | 0.04349 | 0.00034 | Jan-67 | 0 | 0.00034 |
| hc-GIFLP2 | 0.00097 | 0.00686 | Jan-33 | 0 | 0.00034 |
| hic6 | 0.00651 | 0.00034 | 386-1 | 0 | 0.01509 |
| teosin2 | 0.01977 | 0.00686 | hc-C1QL | 0.00097 | 0.00226 |
| hic7 | 0.13207 | 0.00686 | theromacin | 0.00097 | 0.00686 |
| prismafin | 0.00651 | 0.00686 | hc-beta-actin2 | 0.00097 | 0.00034 |
| sovesin | 0.00651 | 0.00686 |  |  |  |

**Figure 3.**
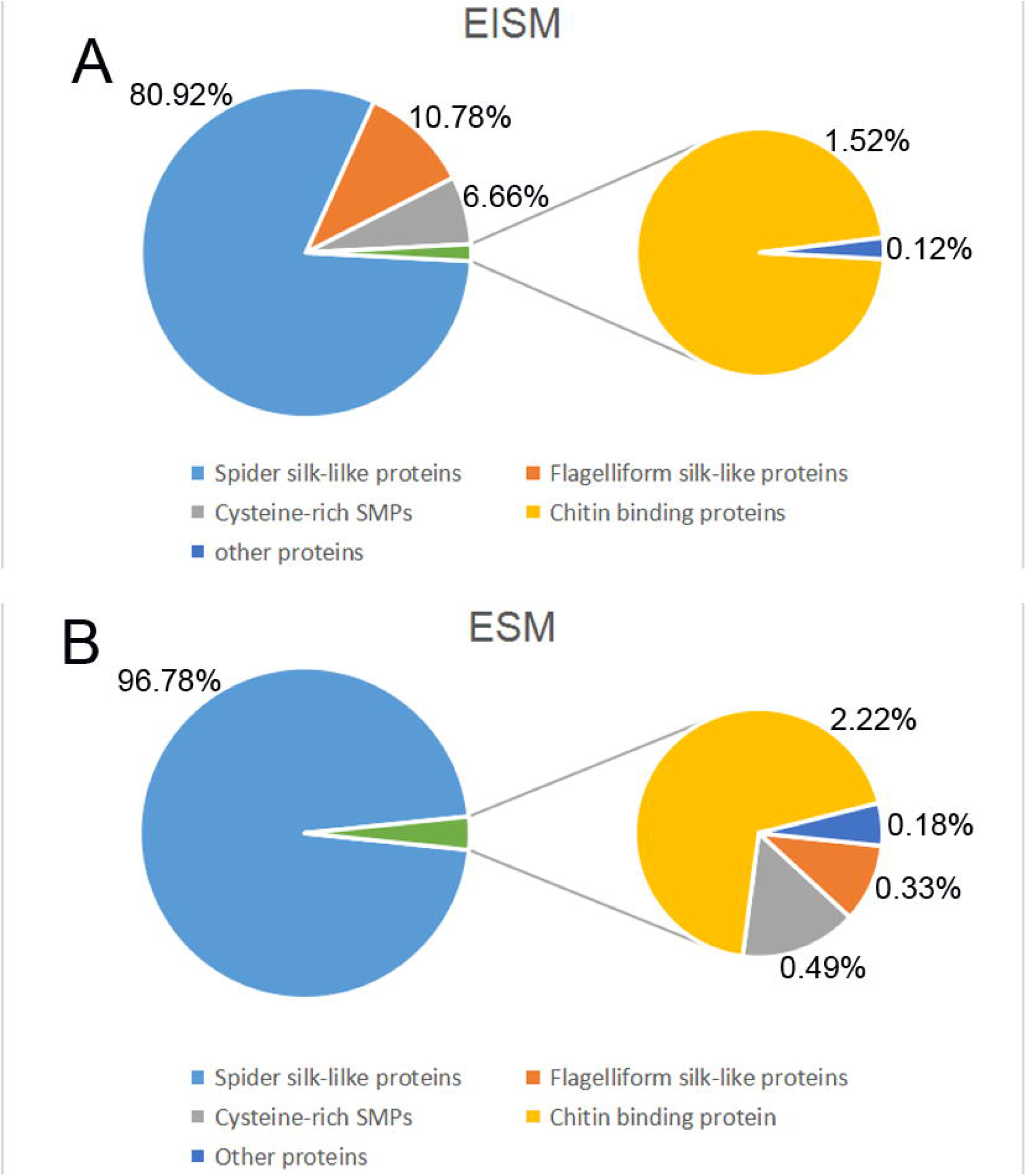
Composition of types of SMPs in EISMs and ESMs. (A) Main SMPs in EISMs were the spider silk-like proteins, flagelliform silk-like proteins, cysteine-rich proteins, and chitin-binding proteins (80.92%, 10.78%, 6.66%, and 1.52% in EISMs, respectively), and the total content of these SMPs was 99.88% in EISMs. The sum content of silk-like proteins (the spider and flagelliform silk-like proteins) was 91.7%. The content of other SMPs was only 0.12%. (B) Main SMPs in ESMs were the spider silk-like proteins and chitin-binding proteins (96.78% and 2.22% in ESMs, respectively), and the total content of these SMPs was 99% in ESMs. The content of the flagelliform silk-like proteins was 0.33%. The sum content of silk-like proteins was 97.11%. Therefore, the content of silk-like proteins in both EISMs and ESMs was more than 90%. As the SMPs were composed of EISMs and ESMs, the content of silk-like proteins in SMPs was more than 90%.

Cytoplasmic and nuclear proteins were complex. Their content was 0.15% in EISMs and 0.02% in ESMs. However, only framework-related proteins (hic68, hic7, hic54, and hc-beta-actin2) were of computational significance, with 0.15% in EISMs and 0.007% in ESMs. Other proteins (hic7-B, hc-RBPL, hc-ZFP385B, hc-NPAC, hc-RBL13, hc-GTLL, hc-EF1A, and hc-ETIF3) were only found in ESMs and occurred in trace amounts (Table 4). The immunity system and protective proteins (cysrichin-D, hic15, hic51, theromacin, hc-C1QL, Krichin, hc-antigen, HcKuPI, prismafin, and hc-GIFLP2) might play an important role in shell protection, but they were more dominant in ESMs (0.07%) than in EISMs (0.02%). The duplication in the aforementioned calculation existed because some proteins contained multiple domains. For example, hic54, hic68 and hic118 were found in both ECM and cytoplasmic and nuclear proteins. “Other proteins” in figure 3 included the cytoplasmic and nuclear proteins and immunity system and protective proteins. The content of “other proteins” in figure 3 did not include these aforementioned multiple domains proteins, because it could not be determined which domains play more important roles. Another reason was that the content of “other proteins” was also very poor even included these aforementioned multiple domains proteins.

## Discussion

A remarkable list of deeply conserved nacre proteins^8,9,20,48^, such as proteins with RLCDs/LCDs (including the spider and flagelliform silk-like proteins), pif, chitin-binding proteins, Kunitz-like proteins, LamG, and so on, was found in the nacre proteome. Some novel or unexpected results were also reported. First, some novel domains of SMPs likely related to energy metabolism were found. QmoC domain (hic7-B) was essential for efficiently delivering electrons to adenosine 5’-phosphosulfate reductase AprAB, which was important in sulfate reduction in sulfate-reducing prokaryotes^49^. Folate_rec domain (hc-RBPL) belonged to folate receptor family, which was attached to the membrane by a Glycosylphosphatidylinositol anchor^50^. Electronic consequences were related to the folate receptor, which bound to folate and reduced folic acid derivatives^51^. Proteins containing these two domains only existed in ESMs. The proteins might be cellular contaminants as well as some cytoplasmic and nuclear proteins^20^. Studies on biomineralization and energy metabolism were few, and most studies focused on bone formation *in vivo*^52^. In shell biomineralization, the processes in mantle, including SMPs synthesis, calcium and hydrogen carbonate transfer, and their secretion, unquestionably need the support of energy metabolism. The relationship between energy metabolism and shell biomineralization *in vitro* remained unknown. However, it also seems unlikely that biomineralization *in vitro*, as a life process, requires no energy at all. The key problem was whether the matrix was preassembled in the mantle cell before secretion because the assembly among different kinds of organic molecules might need energy after they were correctly self-assembled.

Second, protein-synthesis-related proteins might exist systematically in shells. These proteins included protein-synthesis-site-related proteins: such as ribosomal protein (hc-RBL13), glycyl-tRNA synthetase (hc-GTLL), NAC domain-containing protein (hc-NPAC), translation initiation factor (hc-ETIF3), translation elongation factor (hc-EF1A), and transcription activators, such as Med15 domain-containing protein (hic118). hc-GTLL was detected in nacre proteome possibly because glycine was the most dominant amino acid in SMPs^24,25^. The NAC, a complex conserved from archaea to humans, plays an important role in the co-translational targeting of nascent polypeptides to the endoplasmic reticulum^53^. Med15 domain belongs to the sterol regulatory element-binding protein (SREBP) family of transcription activators. SREBPs are critical regulators of cholesterol and fatty acid homeostasis using the evolutionarily conserved ARC105 (also called MED15) subunit to activate target genes^54^. All these proteins were only found in ESMs except hic118, which was only found in EISMs. Cytoplasmic and nuclear proteins were found in almost most of the shell proteome and suspected to be cellular contaminants^20^. This result might be powerful evidence for this opinion.

The last unexpected and the most important phenomenon in this report was that spider silk-like proteins were dominant both in EISMs and ESMs. Two kinds of silk-like proteins exist in nacre: the spider silk-like proteins similar to MIS60 and the flagelliform silk-like proteins similar to MSI30 and shematrin family. They were believed to be insoluble SMPs existing in the organic frameworks^7,37^. The flagelliform silk-like proteins in this study were consistent with this hypothesis; they mainly existed in nacre EISMs (10.78%) and less in ESMs (0.33%) (Figure 3). The spider silk-like proteins (hic74 family, hic58 family, and hic42) were abundant not only in EISMs (80.92%) but also in ESMs (96.78%) (Figure 3). Since the content of silk-like proteins in both EISMs and ESMs was 91.7% and 97.11%, respectively (the sum of the spider and flagelliform silk-like protein contents), we assumed that the content of silk-like proteins in nacre total SMPs exceeded 90%. These results might indicate that the flagelliform silk-like proteins were mainly self-assembled in the nacre organic frameworks, and self-assembled and free state (not be assembled with other organic constituents) spider silk-like matrix proteins states coexisted in nacre. The existence of hic74 and hic58 in the soluble SMPs was also observed in the previous nacre and pearl proteomes of *H. cumingii*^17^. In the first report, hic74 was detected with 17 residues in length peptide “RAAAAAAAAAAAAAIRN” and hic58 was detected in this shell proteome with 28 residues in length peptide “RRFGLDGLGGDSAAAAAAAAAAAGGGRS” and 26 residues in length peptide “RLGSLGGGAAAAAAAAAAAAAGGGRF”^17^. The difference compared with our results was that these proteins were detected only in nacre water and acid-soluble fractions but not in insoluble ones. Therefore, the results from this study provided a new understanding of a previous nacre formation model^26^.

In this model, the first stage of nacre formation was matrix assembly, including the secretion of chitin and alignment by mantle cells; then, other released matrix components were self-assembled to the chitin-binding sites to form ordered organic frameworks^26^. This study indicated that insoluble proteins mainly included silk-like proteins (91.7%), chitin-binding proteins (1.52%), and cysteine-rich SMPs (6.66%) (Figure 3A). The total content of these three kinds of SMPs was 99.88% in EISMs, indicating that these three proteins were the important components of nacre organic frameworks. Chitin-binding proteins were first bound to chitin-binding sites. The content of chitin-related enzymes (chitinase-4 and hc-CDIC) in EISMs (both 0.04349%) was higher than that in ESMs (both 0.00034%) (Table 4), indicating that enzymes might participate in the arrangement of chitins by modifying them. The role of cysteine-rich SMPs might be more important than anticipated, as their higher content in EISMs (6.66%) was far more than that in ESMs (0.49%) (Figure 3). Cysteine participated in protein interaction through disulfide bond formation. All chitin-binding proteins had rich cysteine residues. MSI60^7^, hic74 family, and hic42 all have cysteine residues in their N-terminal regions except hic58 family. hic58-3 had two amino acid mutations from “GS” to “CT” at 198 and 199 sites (Figure S11), and its content (12.96%) was higher than those of hic58-1 (0.88%) and hic58-2 (0.08%) in EISMs (Table 4), although hic58-3 had only a cysteine residue after mutation. This result might indicate the importance of disulfide bonds in the organic framework self-assembly. In flagelliform silk-like proteins, silkmapin, silkmaxin, and hic9 had no cysteine residues, but hic10 and metrichin had two and six cysteine residues, respectively. Therefore, how silkmapin, silkmaxin, hic9, hic58-1, and hic58-3 were assembled into organic frameworks might need in-depth investigation. Cysteine-rich proteins might act as “connectors”. Based on these results, chitin assembly was assumed to have been helped by SMPs: (1) chitin-binding proteins first bound to chitin in the first step; (2) silk-like proteins bound to chitin-binding proteins themselves or were helped by cysteine-rich proteins to form network structures; and (3) cysteine-rich proteins might act as the connectors and fixers to facilitate the binding of the chitin-binding proteins, silk-like proteins, and themselves with chitin.

The second step of the matrix assembly in the model was space filling with the silk gel in the organic frameworks^26^. This assumption was well supported by this study because 97.11% of ESMs were silk-like proteins. This study also indicated that 96.78% of ESMs were spider silk-like proteins and only 0.33% were flagelliform silk-like proteins (Figure 3B); therefore, the silk gel was mainly composed of spider silk-like proteins. Spider silk-like proteins of nacre SMPs, such as MSI60, hic74 family, hic58 family, and hic42, constitute the (poly-A or A-rich or poly-GA)-(poly-G or G rich) repeated blocks (the number of poly-GA block was always 1), and hydrophilic amino acids existed in G-rich blocks (Figure 4). This resulted in spider silk-like proteins having an alternate permutation of hydrophilic and hydrophobic blocks in their amino acid sequence similar to the spider silk protein (Figure 4). Spider silk proteins were always self-folding to be globular proteins named “fibroin micelle,” following the assembly stage to form fibroin globule using fibroin micelles as basic molecules, and these fibroin globule could be aligned and assembled continually^55^. The alternate permutation of hydrophilic and hydrophobic blocks enabled the hydrophobic blocks to be assembled and organized into micelles depending on chain folding and hydrophobic interactions, with the hydrophilic blocks on the micelle surface; they remained in gel state until fibroin globule was formed^55^. Alternative splicing should also be considered in the SMP self-assembly, as they might generate silk-like proteins of different sizes. This needs further exploration. Therefore, spider silk-like proteins in nacre might form irregular-sized micelles depending on a similar folding mechanism. According to the model of the spider silk protein assembly, the hydrophilic blocks promoted the solubility of micelles in water^55^.

**Figure 4.**
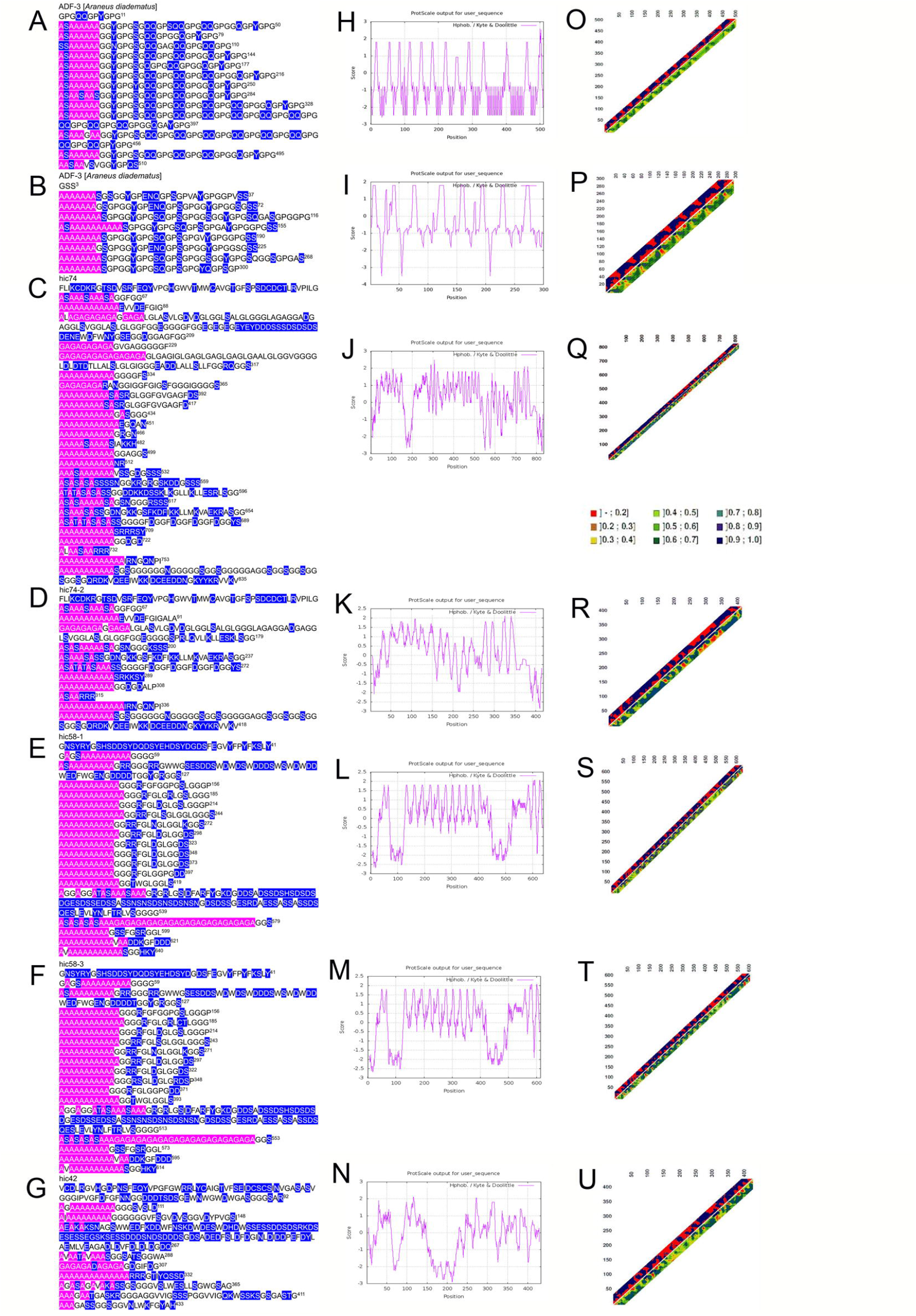
RLCDs of partial spider fibroin proteins (ADF-3 and ADF-4) and nacre spider silk-like SMPs (hic74, hic74-2, hic58-1, hic58-3, and hic42) and their hydrophobicity and protein distance analysis. (A–G) RLCDs of ADF-3, ADF-4, hic74, hic74-2, hic58-1, hic58-3, and hic42. The amino acids in the pink box were poly-A, poly-GA-repeat, and A-rich blocks. The amino acids in the blue box were hydrophilic amino acids. (H–N) Hydrophobicity analysis of ADF-3, ADF-4, hic74, hic74-2, hic58-1, hic58-3, and hic42. The hydrophilic and hydrophobic blocks were alternately permuted in partial spider fibroin proteins and nacre spider silk-like SMPs according to (A–N). (O–U) The protein distance constraint analysis of ADF-3, ADF-4, hic74, hic74-2, hic58-1, hic58-3, and hic42. The protein distance constraint analysis could investigate the correlations between sequence separation (in residues) and distance (in Angstrom) of any pair of amino acids in polypeptide chains. The results were only for reference to show the similar assembly of partial spider fibroin proteins and nacre spider silk-like SMPs. The upper triangle denotes the distance constraints of published coordinates, where red points indicate distances above the thresholds (mean values) and dark points depict distances below the threshold. The lower triangle had no significance. The poly-A and poly-GA-repeat should be in the interior of the protein structure similar to partial spider fibroin proteins.

It was possible that the solubility and insolubility of a protein in the extraction from the shell were determined not only by their nature but by the level of assembly with other proteins. Especially when intermolecular covalent bonds were formed, the soluble proteins might occur in insoluble SMPs with other framework proteins. A similar phenomenon existed in chitin-binding proteins, which were another important soluble constituent with a content of 2.22% in ESMs. hic124 and hc-pif2 were both dominant in EISMs and ESMs, and hichin was more dominant in ESM (0.1%) than in EISM (0.00097%). pif played dual key roles in nacre formation by binding to chitin and regulating aragonites crystal growth direction^56^. The results of this study were consistent with the roles of pif. Chitin-binding proteins and silk-like proteins accounted for 99.33% of ESMs. Although the importance of SMPs in biomineralization was not determined by their quantities, the dominant SMPs should be considered to form the microenvironment of calcium carbonate crystallization. As the microspace of organic frameworks was filled with the silk gel composed mainly of spider silk-like proteins in nacre, they inevitably interacted with calcium carbonate. The green mussel *Perna canaliculus* shells revealed 2-to 25-nm intracrystalline organic inclusions in the nacreous layer using High angle annular dark field image-STEM; no organic agglomerates were found in the area of 50 nm adjacent to the interlamellar organic sheets^57^. A relation might exist between “fibroin micelles” mentioned above and the intracrystalline organic inclusions in nacre of *P. canaliculus* as spider silk-like proteins accounted for 96.78% of ESMs, but deeper studies are needed for direct evidence. A recombination green fluorescent protein labeled abalone SMP lectin (GFP-Lectin) was used to conduct calcium carbonate crystallization experiments *in vitro*. GFP-Perlucin was confirmed to be encapsulated into the crystal lattice^58^. Then, Pendola and Evans proposed that the physical properties of the nacre matrix protein hydrogels dictated hydrogel–ion interactions^59^. Spider silk-like proteins might participate in calcium carbonate crystallization, as spider silk-like proteins were acidic and had a concentrated distribution of acidic amino acids except for hic74-2, with 0.07% content in ESMs. Therefore, spider silk-like proteins might have three roles in nacre biomineralization: (1) silk-like proteins were the main protein constituents of organic frameworks and participated in organic framework formation together with chitin-binding proteins and cysteine-rich proteins; (2) free-state spider silk-like proteins formed a silk gel to fill the microspace of organic frameworks; and (3) silk-like proteins also participated in calcium carbonate crystallization with other SMPs.

## Methods

### Shell preparation and protein extraction

Shell preparation and protein extraction were performed by the methods described in a previous study^18^. The shells of *H. cumingii* were cleaned and immersed in 5% sodium hydroxide for 24 h to move possible organic pollution on the inner surface of the shell nacreous layer. After cleaning with distilled water, the prismatic layer was moved by abrasion before air-drying. Nacre powders were decalcified with 0.8 M EDTA to obtain ESMs. The insoluble organic matrix was cleaned after complete decalcification; then, EISMs were obtained by the method described in a previous study^18^. All processes were performed under sterile conditions.

### Label-free proteomic analysis

The proteins of nacre ESMs and EISMs were replaced three times with 8M urea, and the concentration was determined by the Bradford method. Sodium dodecyl sulfate–polyacrylamide gel electrophoresis (SDS-PAGE) electrophoresis was used to analyze the protein samples and evaluate whether the quality of the samples met the requirements of subsequent experiments (Figure 1S). Then, 200 μL of protein samples were mixed with TCEP to a final concentration of 10mM and reacted for 60 min at 37 °C. Then, iodoacetamide was added to a final concentration of 40mM and reacted at room temperature for 40 min in the dark. Precooled acetone was added to each tube (acetone:sample volume ratio = 6:1) and precipitated for 4 h at −20 °C. The precipitate was collected after centrifuging at 10,000 *g* for 20 min and dissolved thoroughly in 200 μL of 100mM tetraethylammonium bromide. Trypsin was added to each tube at a mass ratio of 1:50 (enzyme:protein) and enzymolyzed overnight at 37 °C. After trypsin digestion, peptides were drained with a vacuum pump and redissolved in 2% ACN and 0.1% TFA.

After desalting, the peptide was drained with a vacuum concentrator and dissolved in mass spectrometry loading buffer to 0.5 μg/μL for liquid chromatography with tandem mass spectrometry (MS) analysis using a Q-Exactive mass spectrometer (Thermo, USA). Four samples with three replicates per sample were used (ESMs from white nacre: WS1, WS2, and WS3; ESMs from purple nacre: PS1, PS2, and PS3; EISMs from white nacre: WI1, WI2, and WI3; EISMs from purple nacre: PI1, PI2, and PI3). C18 column (Thermo, USA) was used for MS analysis, and Thermo Xcalibur 4.0 (Thermo, USA) was used for data acquisition. The MS scanning range (*m*/*z*) was 350–1300, with the acquisition mode of data-dependent acquisition. The Primary MS resolution was 70,000, with the fragmentation mode of High-energyC-trap Dissociation. Secondary MS resolution was 17,500, with a dynamic exclusion time of 30 s. An established database was selected, and the raw file was submitted to the server for database search. The project database website was https://www.ncbi.nlm.nih.gov/protein. The filter parameter was peptide FDR ≤ 0.01.

### RACE and bioinformatics analysis

Based on the EST sequences matched with peptides of unknown SMPs, primers were designed for 5’ and 3’ RACE polymerase chain reactions (Table S1-2). The proteins with full-length cDNA sequence were analyzed with the following programs: The ORF was detected with the ORF finder (http://www.ncbi.nlm.nih.gov/projects/gorf/). The protein domain prediction homology analysis was performed using GenBank and BLAST. The signal peptide was detected using the SignalIP4.1 Server (http://www.cbs.dtu.dk/services/SignalP/). The amino acid sequence of the mature protein was analyzed using ExPasy (http://web.expasy.org/protparam/). The hydrophobicity of proteins was analyzed using ProtScale (https://web.expasy.org/protscale/). The protein distance constraints were analyzed using distanceP (http://www.cbs.dtu.dk/services/distanceP/). The protein concentration was calculated by the method developed by States et al^45^.

## Acknowledge

This work was financially supported by the National Natural Science Foundation of China (32072975).

