## Supplemental Figure 1-15 for "More than 90% of nacre matrix proteins are composed of silk-like proteins": Supplied Materials Online.docx

**This PDF includes:**

Figures.S1 to S15

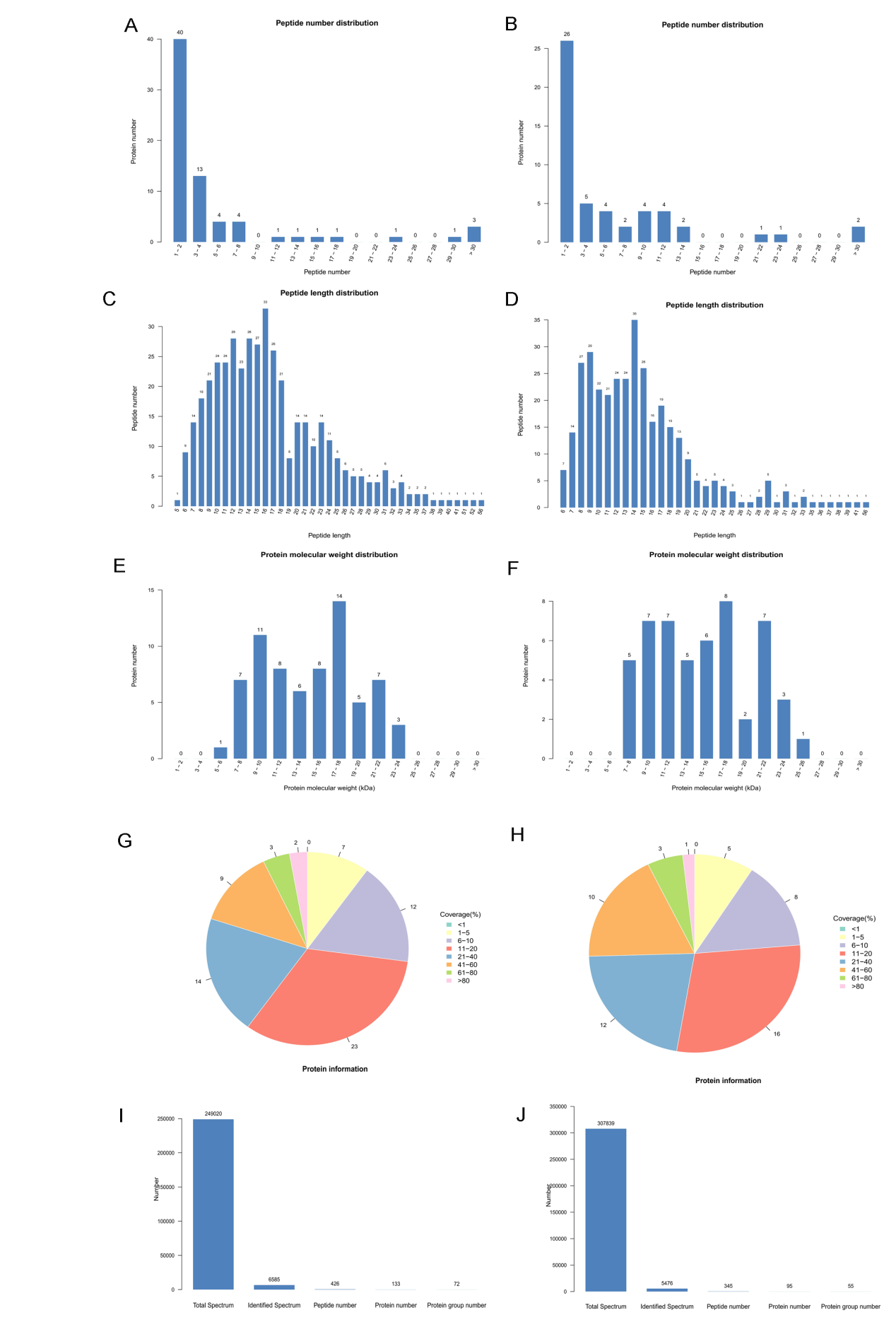

Figure S1. Quality assessment of the nacreous proteome. A, peptide number distribution of nacre ESMs; B, peptide number distribution of nacre EISMs; C, peptide length distribution of nacre ESMs; D, peptide length distribution of nacre EISMs; E, protein molecular weight distribution of nacre ESMs; F, protein molecular weight distribution of nacre EISMs; G, Protein coverage distribution of nacre ESMs; H, Protein coverage distribution of nacre EISMs; I, identified protein information of nacre ESMs; J, identified protein information of nacre EISMs.

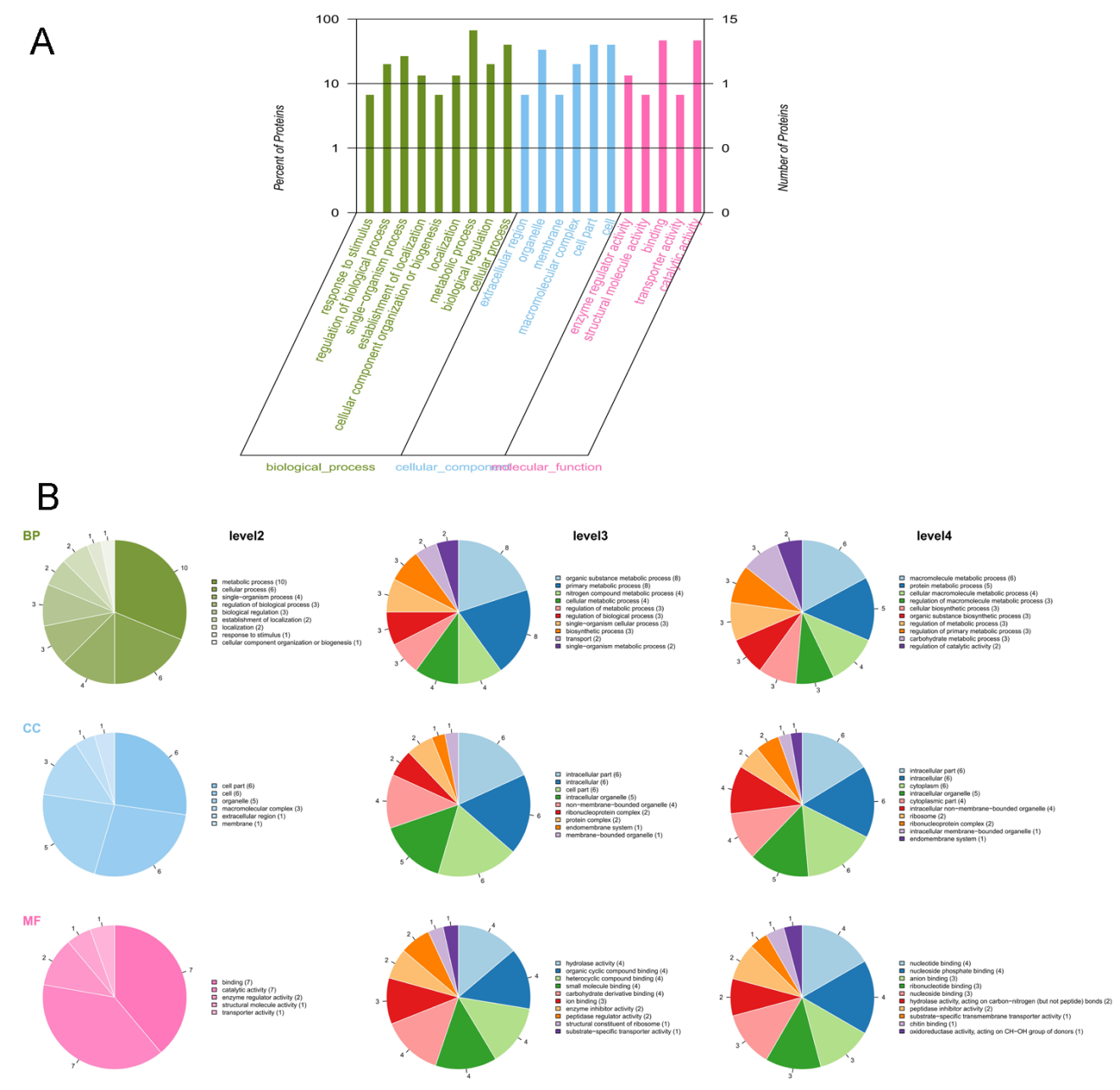

Figure S2. GO (Gene Ontology) function classification annotation. A, GO secondary classification statistics (Each column in the figure represents the secondary classification of a GO, and the higher the column, the more proteins in this secondary classification. The abscissa represents the secondary classification terms of GO; The ordinate (left) represents the percentage of proteins included in the secondary classification, and the ordinate (right) represents the number of proteins in the secondary classification; The three colors represent the three major groups, with green representing biological processes, blue representing cellular components, and red representing molecular functions) ; B, GO 2, 3, 4 level classification statistics chart (Different colors in each pie chart in the figure represent different GO terms, and its area represents the relative proportion of proteins in the GO Term. This figure contains 9 pie charts, with 3 rows from top to bottom, representing the three branches of GO, namely BP (biological process, green cake), CC (cell component, blue cake) and MF (molecular function, pink cake). From left to right, there are three columns, which respectively represent the more detailed classification under the three GO branches, namely, level2, 3, and 4. As the number of layers increases, the descriptions of BP, CC, and MF become more detailed, and the functions of the protein from level2 -> level3 -> level4 are more clearly annotated).

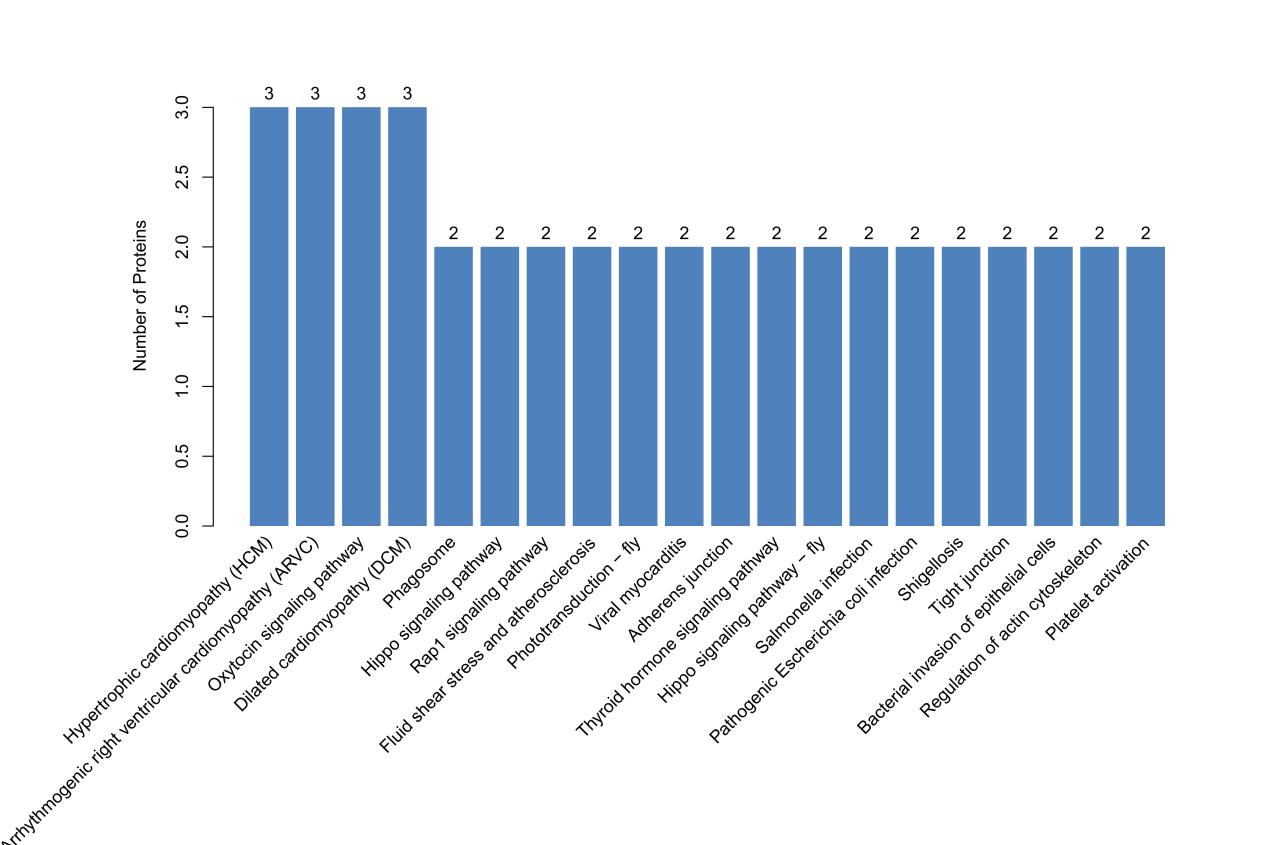

Figure S3. Top 20 pathways with the largest number of proteins.

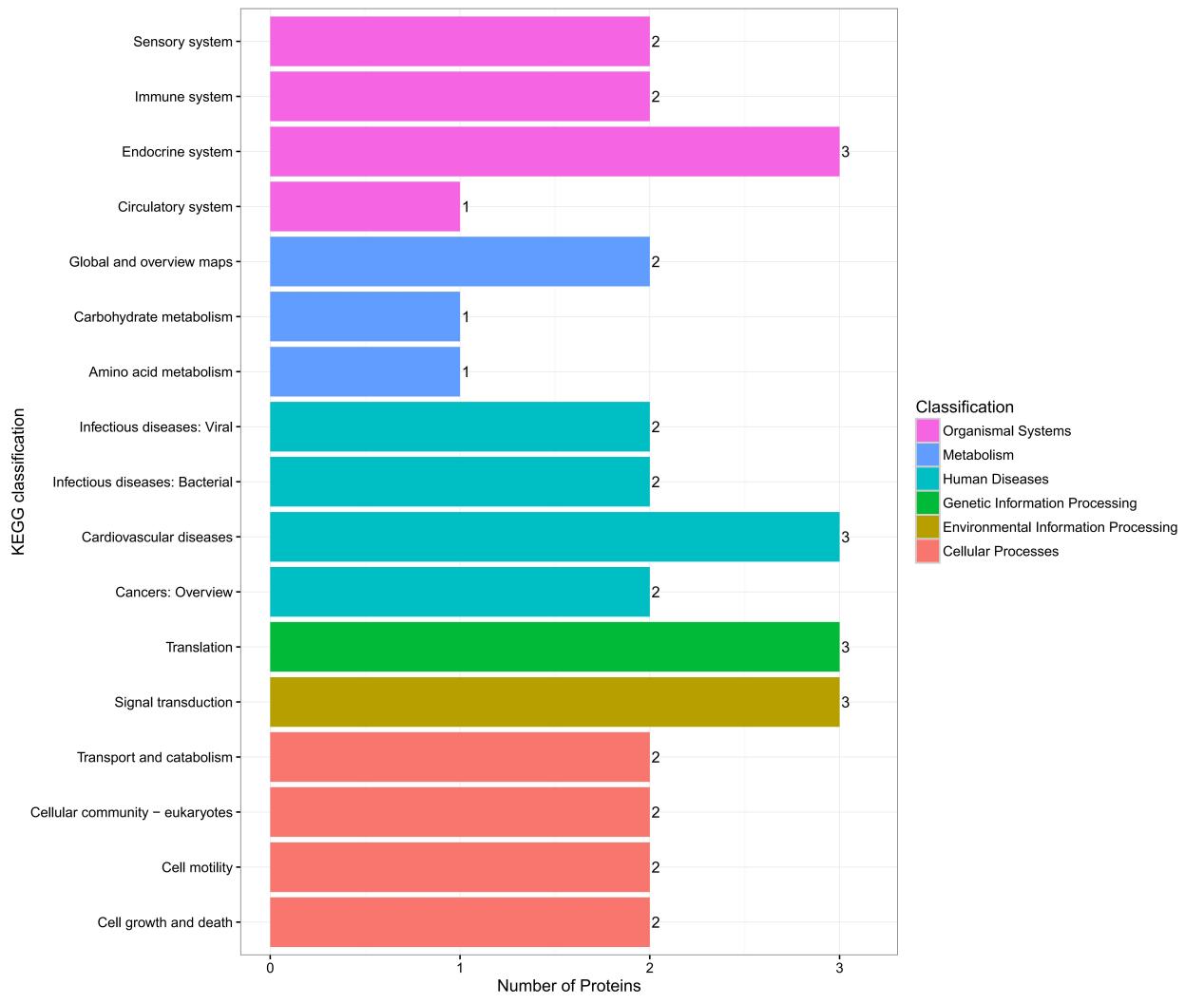

Figure S4. KEGG annotation statistics. The ordinate is the name of the KEGG metabolic pathway, and the abscissa is the number of proteins annotated to the pathway.

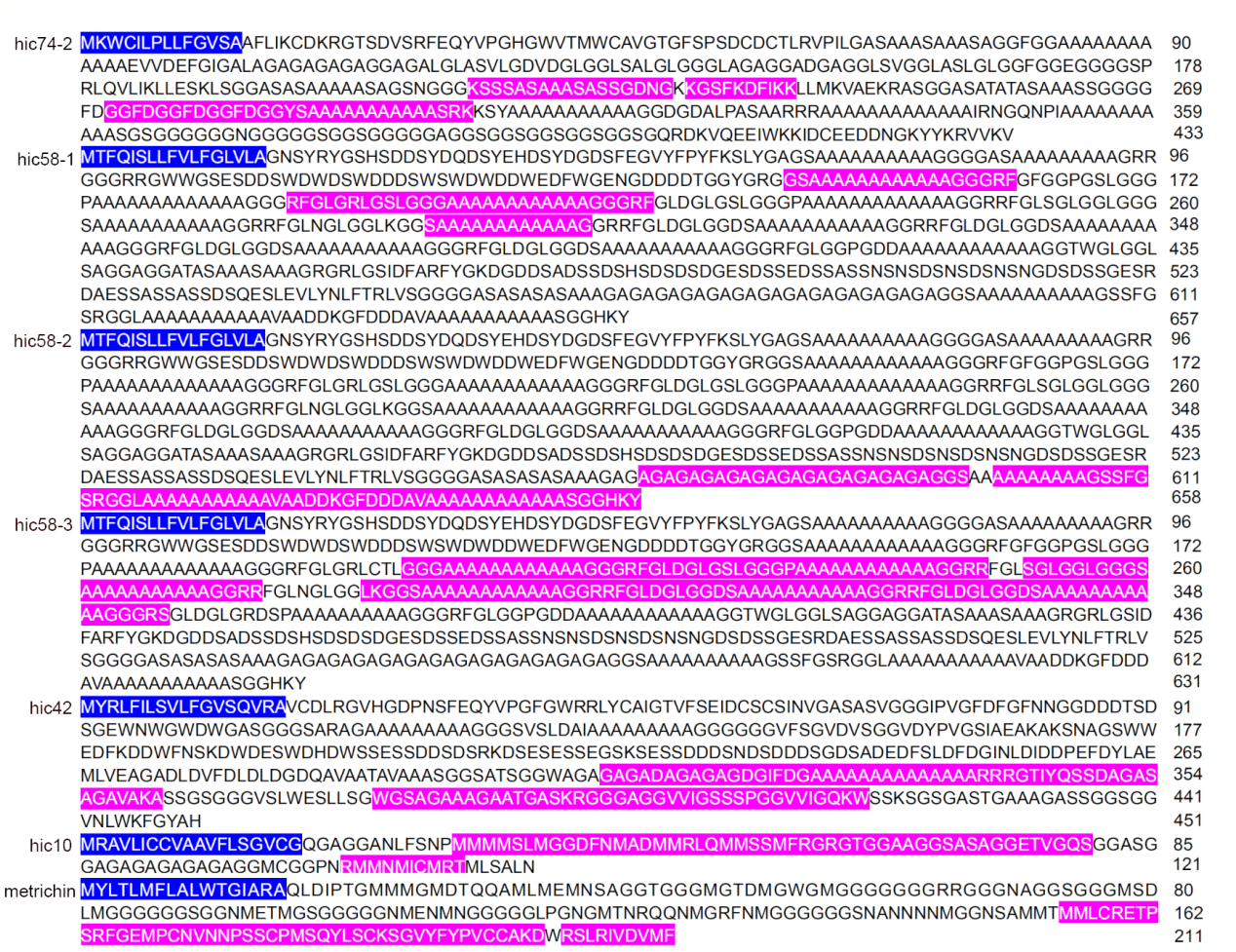

Figure S5. Complete amino acid sequences of hic74-2, hic58-1, hic58-2, hic58-3, hic42, hic10 and metrichin. The cDNA sequences of there proteins was submitted to Genebank (hic74-2, Accession No. MZ422369; hic58-1 to hic58-3, Accession No. MZ422366-MZ422368; hic42, Accession No. MZ422370; hic10, Accession No. MZ422371; metrichin, Accession No. MZ440742).The amino acid sequences of every protein in the blue box were signal peptides. The amino acid sequences of every protein in the pink box were detected in nacre matrix protein proteome.

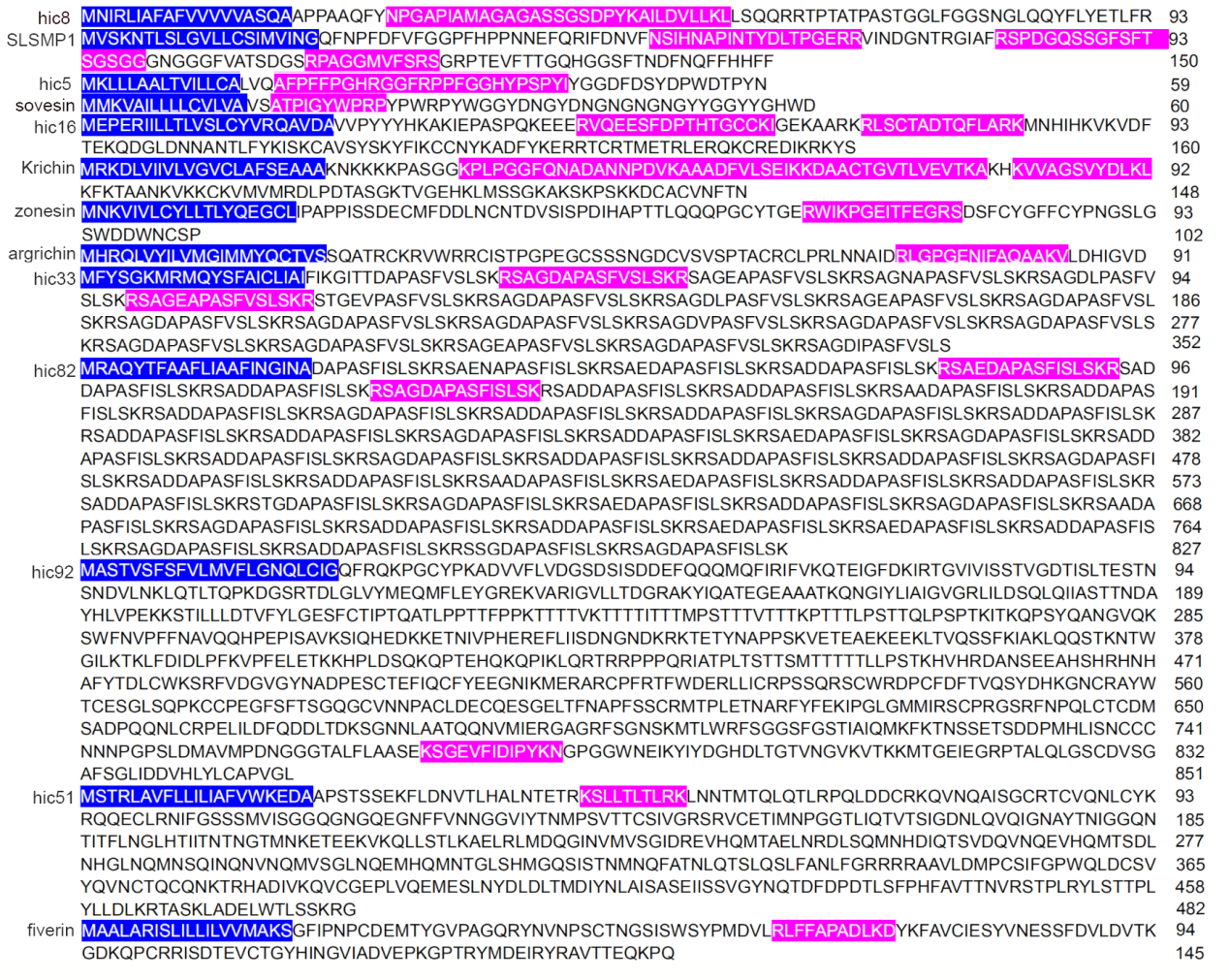

Figure S6. Complete amino acid sequences of hic8, SLSMP1, hic5, sovesin, hic16, Krichin, zonesin, argrichin, hic33, hic82, hic92, hic51 and fiverin. The cDNA sequences of there proteins was submitted to Genebank (hic8, Accession No. MZ422372; SLSMP1, Accession No. MN829945; hic5, Accession No. MZ440754; sovesin, Accession No. MZ440743; hic16, Accession No. MZ467108; Krichin, Accession No. MF371330; zonesin, Accession No. MZ467100; argrichin, Accession No. MZ440755; hic33, Accession No. MZ440758; hic82, Accession No. MZ440759; hic92, Accession No. MZ440757; hic51, Accession No. MZ467109; fiverin, Accession No. MZ467101). The amino acid sequences of every protein in the blue box were signal peptides. The amino acid sequences of each protein in the pink box were detected in nacre matrix protein proteome.

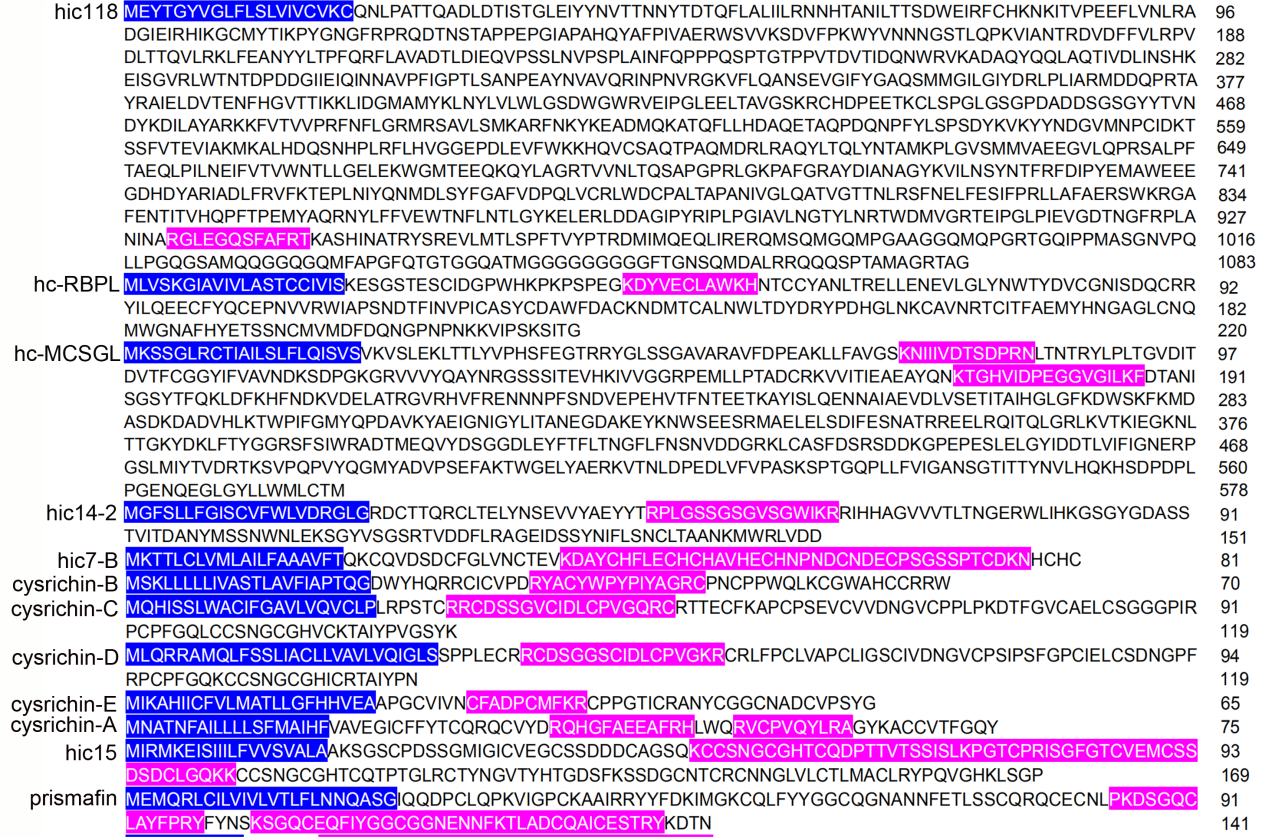

Figure S7. Complete amino acid sequences of hic118, hc-RBPL, hc-MCSGL, hic14-2, hic7-B, cysrichin-B, cysrichin-C, cysrichin-D, cysrichin-E, cysrichin-A, hic15 and prismafin. The cDNA sequences of there proteins was submitted to Genebank (hic118, Accession No. MZ440756; hc-RBPL and hc-MCSGL, Accession No. MZ467110 and MZ467111; hic14-2 and hic7-B, Accession No. MZ467103 and MZ467104; cysrichin-B, Accession No. MZ440752; cysrichin-C, cysrichin-D, and cysrichin-E, Accession No. MZ467105, MZ467106 and MZ467107; cysrichin-A, Accession No. MZ440751; hic15, Accession No. MN829944; prismafin, Accession No. MN829947). The amino acid sequences of every protein in the blue box were signal peptides. The amino acid sequences of each protein in the pink box were detected in nacre matrix protein proteome.

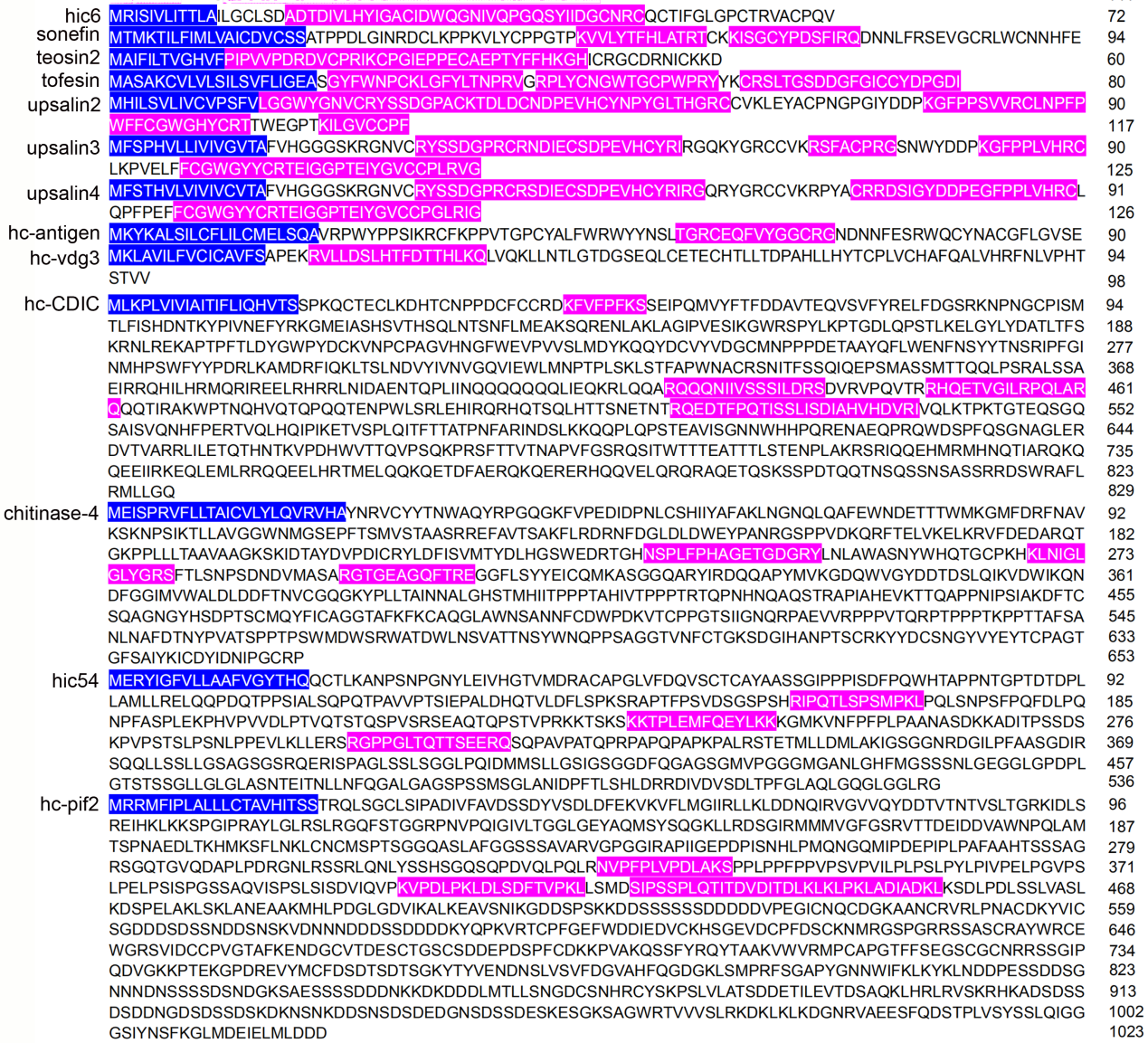

Figure S8. Complete amino acid sequences of hic6, sonefin, teosin2, tofesin, upsalin2, upsalin3, upsalin4, hc-antigen, hc-vdg3, hc-CDIC, chitinase-4, hic54 and hc-pif2. The cDNA sequences of there proteins was submitted to Genebank (hic6, Accession No. MG722969; sonefin, Accession No. MZ467099; teosin2, Accession No. MZ467098; tofesin, Accession No. MZ440747; upsalin2, upsalin3 and upsalin4, Accession No. MZ440748, MZ440749 and MZ440750; hc-antigen, Accession No. MZ467102; hc-vdg3, Accession No. MZ467112; hc-CDIC, Accession No. MZ467097; chitinase-4, Accession No. MZ467096; hic54, Accession No. MZ440745; hc-pif2, Accession No. MZ440746). The amino acid sequences of every protein in the blue box were signal peptides. The amino acid sequences of each protein in the pink box were detected in nacre matrix protein proteome.

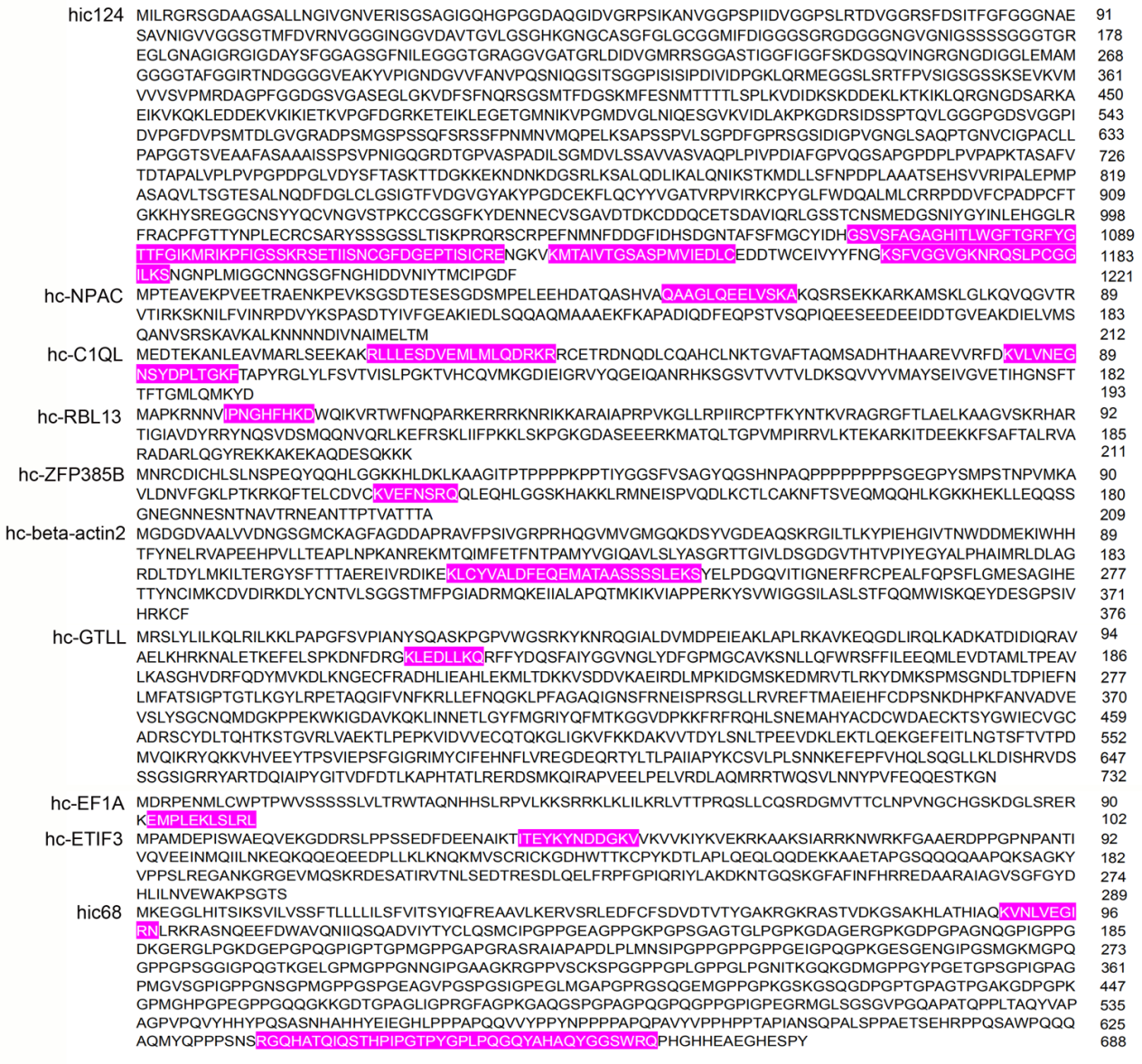

Figure S9. Complete amino acid sequences of hic124, hc-NPAC, hc-C1QL, hc-RBL13, hc-ZFP385B, hc-beta-actin2, hc-GTLL, hc-EF1A, hc-ETIF3 and hic68. The cDNA sequences of there proteins was submitted to Genebank (hic124, Accession No. MZ440744; hc-NPAC, Accession No. MZ467113; hc-C1QL, Accession No. MZ467119; hc-RBL13, Accession No. MZ467114; hc-ZFP385B, Accession No. MZ467115; hc-beta-actin2, Accession No. MZ467120; hc-GTLL, Accession No. MZ467116; hc-EF1A, Accession No. MZ467117; hc-ETIF3, Accession No. MZ467118; hic68, Accession No. MZ440753). The amino acid sequences of each protein in the pink box were detected in nacre matrix protein proteome.

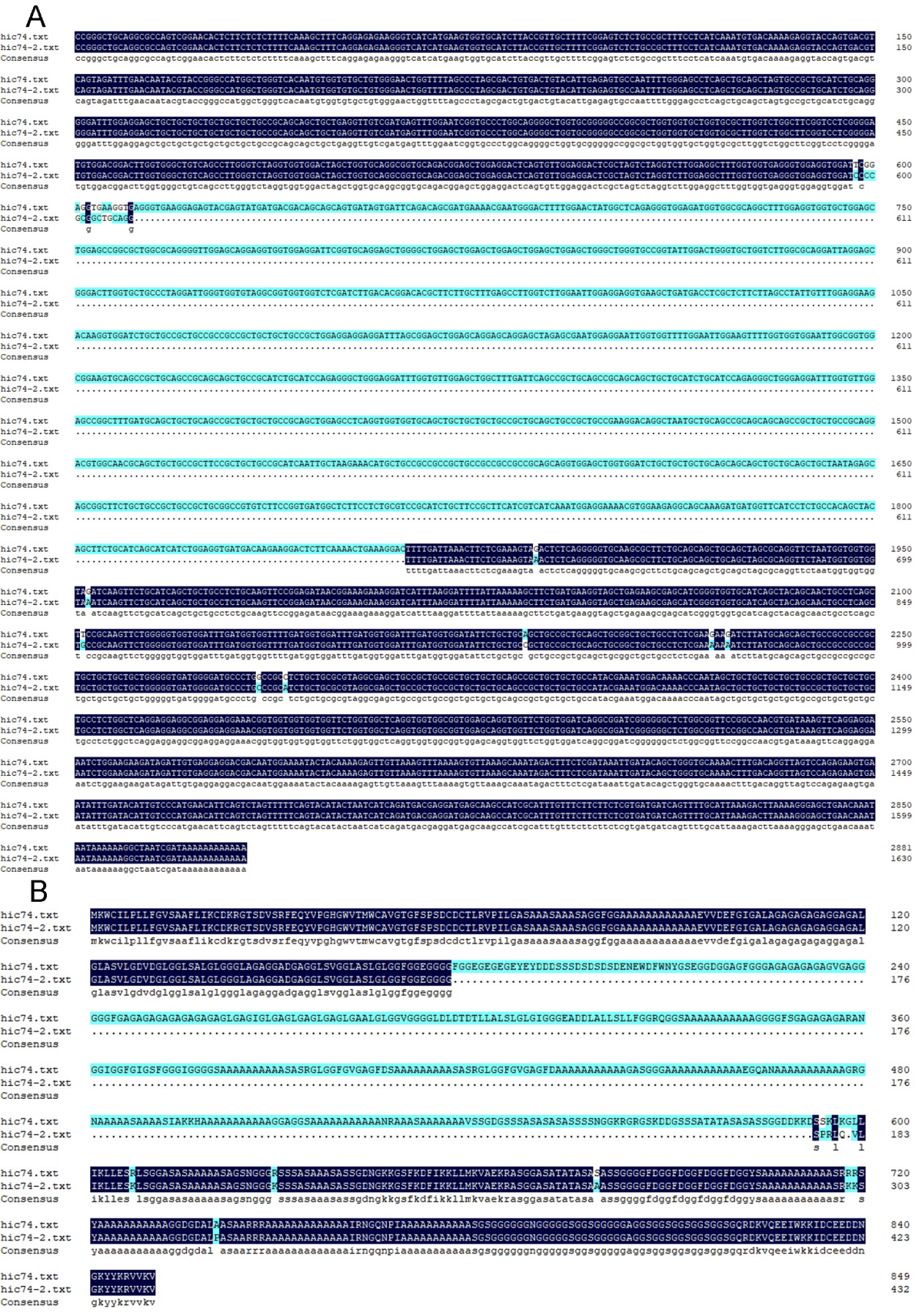

Figure S10. Comparison of the nucleic acid sequences (A) and the predicted amino acid sequences (B) of hic74 and hic74-2.

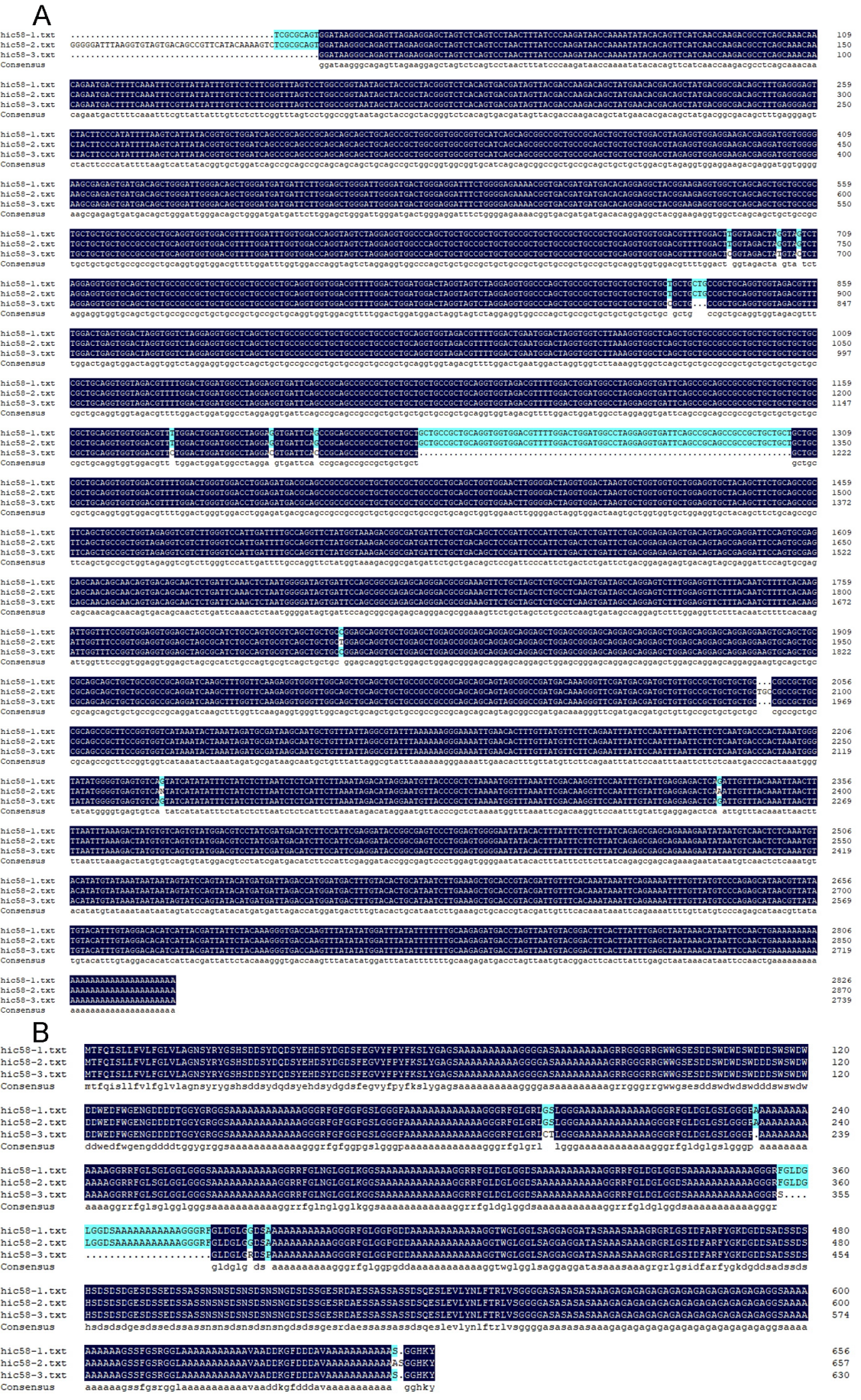

Figure S11. Comparison of the nucleic acid sequences (A) and the predicted amino acid sequences (B) of hic58-1 to -3.

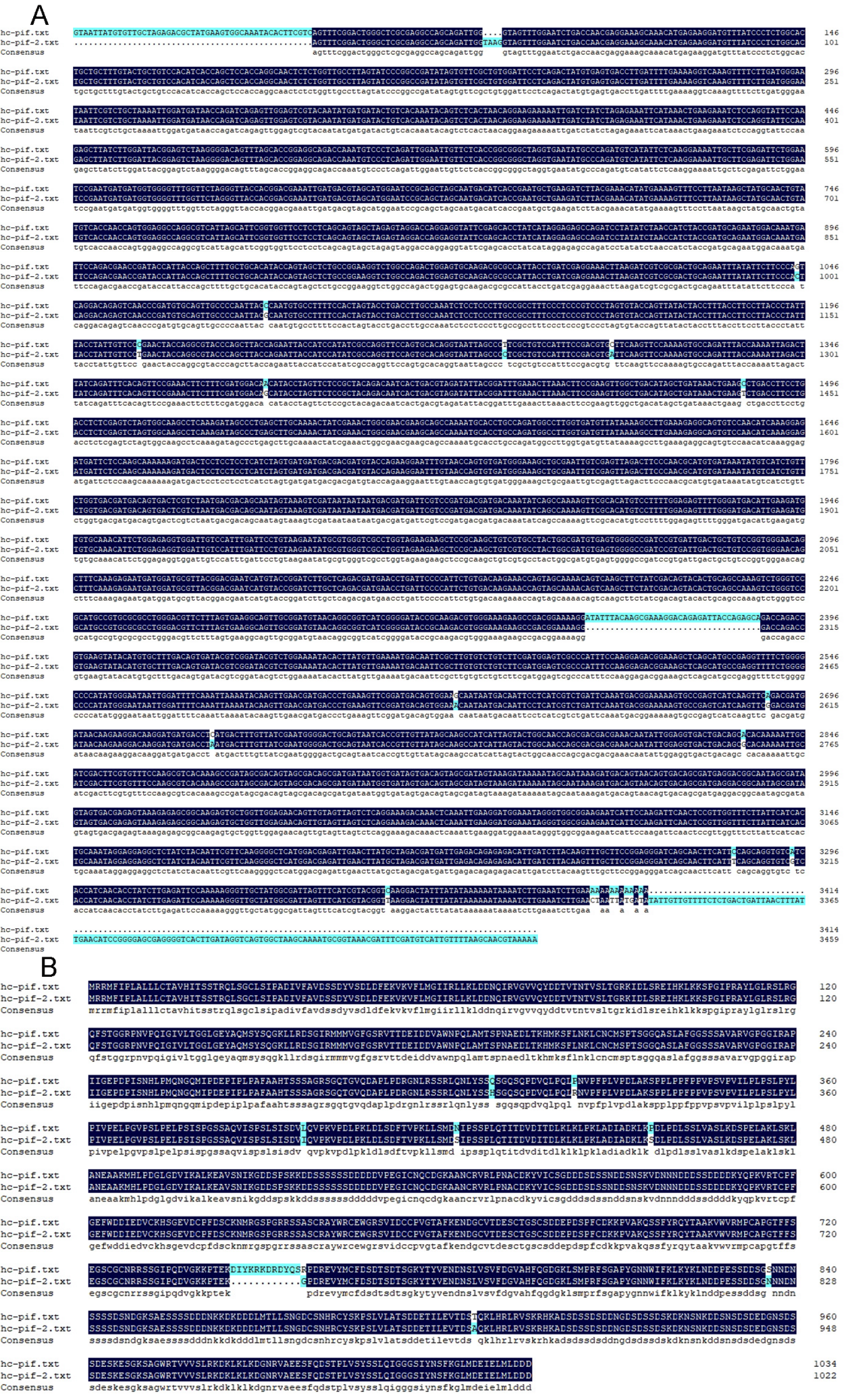

Figure S12. Comparison of the nucleic acid sequences (A) and the predicted amino acid sequences (B) of hc-pif and hc-pif-2.

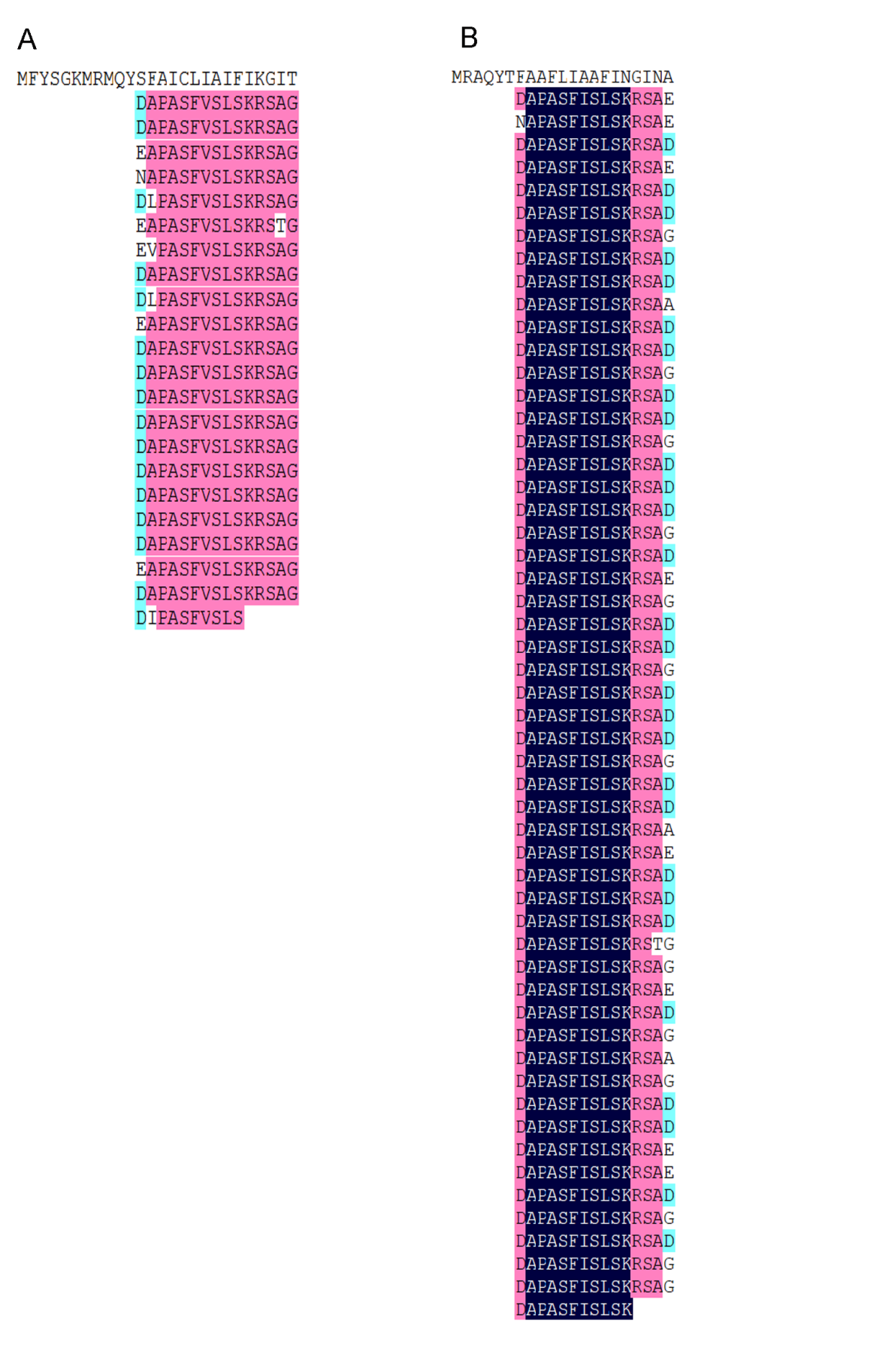

Figure S13. The repeat sequence of hic33 and hic82.

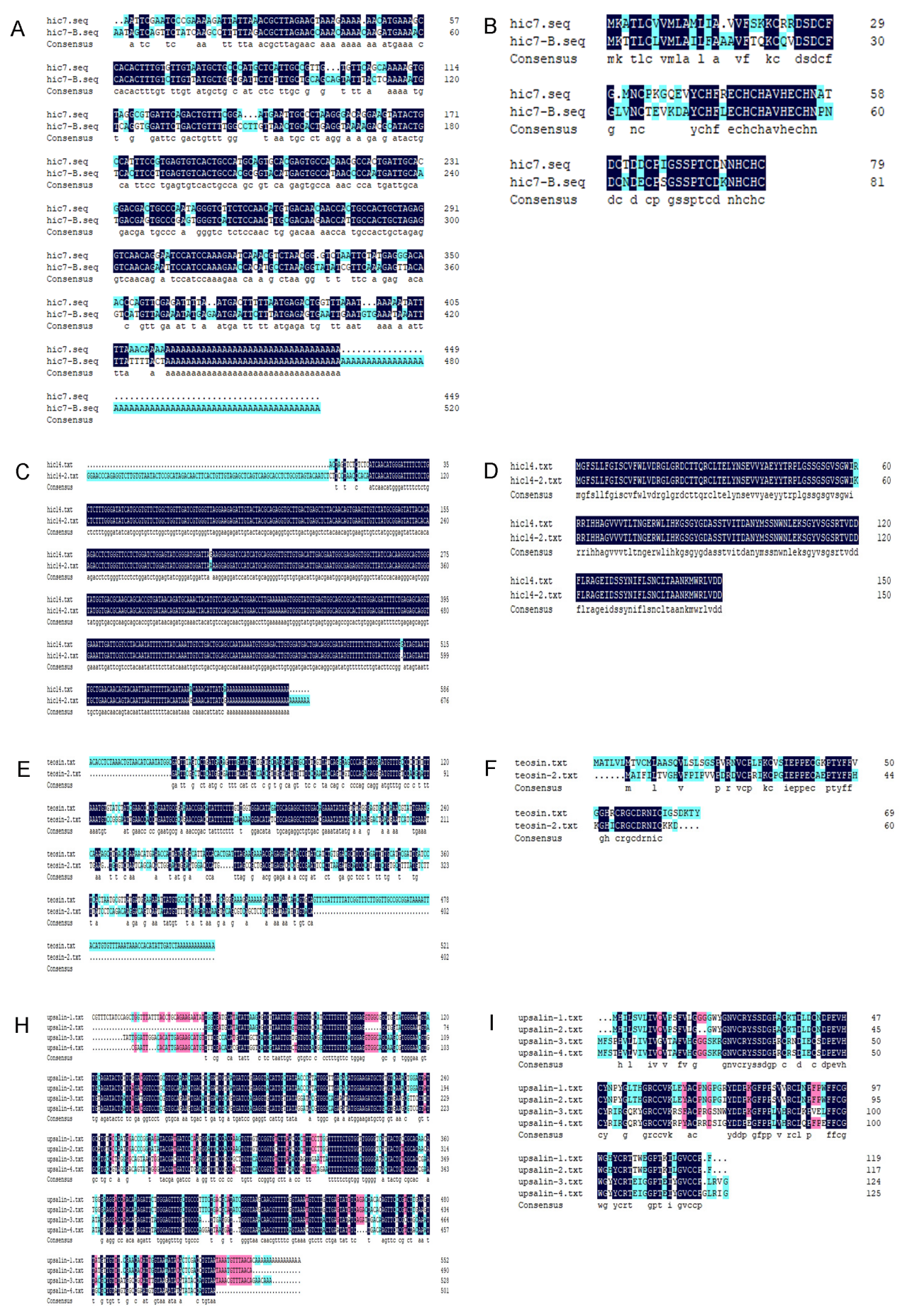

Figure S14. Comparison of the nucleic acid sequences and the predicted amino acid sequences of hic7 and hic7B (A and B), hic14 and -2 (C and D), teosin and -2 (E and F), upsalin family (H and I).

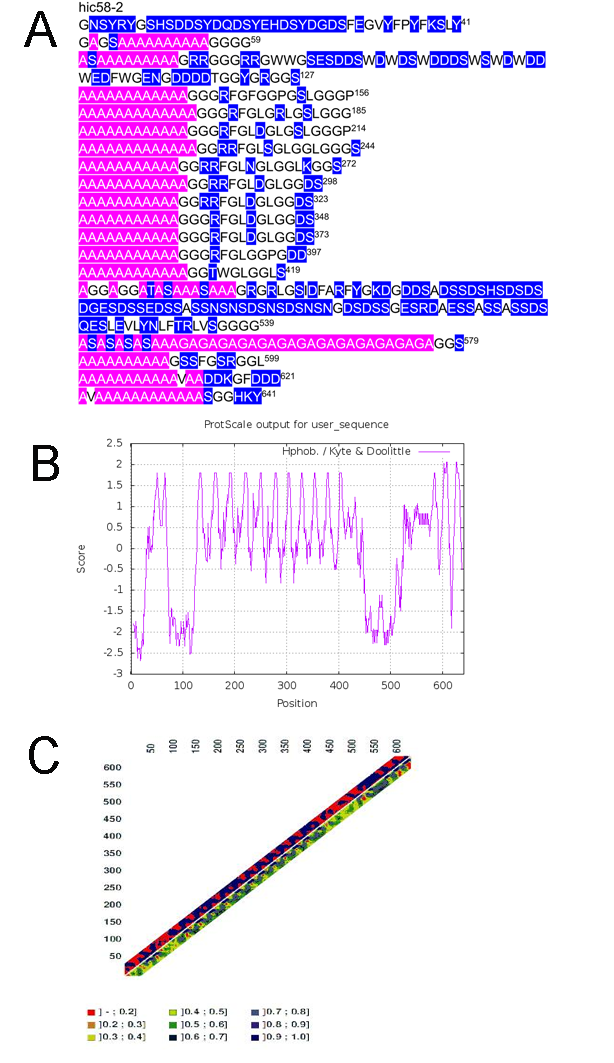

Figure S15. Additional information of hic58-2.
